# Spatially confined dopamine from locus coeruleus axons selectively shapes hippocampal dynamics and action timing

**DOI:** 10.64898/2026.09.02.748730

**Authors:** Dinghao Luo, Jingyu Cao, Raphael Heldman, Lin Tian, Yingxue Wang

## Abstract

Neuromodulatory systems project broadly, yet computations engage specific neuronal populations. Whether neuromodulation can selectively influence task-engaged neurons remains unclear. During goal-directed navigation, neurons in the locus coeruleus (LC), the brain’s principal norepinephrine source, responded at navigation onset. Concurrently, dopamine transients arose in micrometer-scale domains around LC axons in CA1. These transients preferentially enhanced nearby CA1 neurons with ramping dynamics, whose higher activity predicted later reward-anticipatory action. Brief LC activation evoked local dopamine, enhanced ramping dynamics, and delayed action initiation seconds later. Blocking D_1_-like, but not adrenergic, receptors weakened ramping dynamics and impaired reward anticipation. Modeling showed that selectivity can emerge when localized dopamine coincides with strong ongoing neuronal activity, without dedicated wiring. Thus, a widely projecting system can selectively modulate task-engaged neurons and shape behavior.

## Introduction

Neuromodulatory systems regulate arousal, attention and motivation^1–4^. They project widely across the brain, and can influence large populations of neurons^5–7^. Yet ongoing computations often depend on coordinated activity among particular subsets of neurons^8–10^. This mismatch raises a fundamental question: can broadly projecting neuromodulatory systems selectively influence the right neurons at the right time, and if so, by what mechanism?

The locus coeruleus (LC) provides a useful system for addressing this question. This small brainstem nucleus is the brain’s principal source of norepinephrine (NE)^1–3^ and innervates many brain regions, including hippocampal CA1^3,11–15^. LC axons in CA1 show phasic increases in activity around defined behavioral events, including the onset of locomotion^16,17^, raising the possibility that this pathway may modulate hippocampal activity at specific moments. Although the LC is canonically noradrenergic, emerging evidence has also implicated the LC as a source of dopamine (DA) in the dorsal hippocampus^14,15,18,19^. However, it remains unclear whether phasic LC input selectively shapes CA1 neuronal dynamics during an ongoing computation, and whether any such effect is mediated by DA or NE.

To examine these questions, we focused on goal-directed navigation, a computation where CA1 dynamics have been well characterized. During navigation, animals track their position and use this information to guide behavior. CA1 neurons can fire at specific spatial locations^20^, elapsed times^21,22^, or traveled distances^23^, and their activity can also vary according to the action an animal subsequently performs^21,24,25^. Beyond these discrete firing patterns, we recently identified PyrUp neurons, whose activity rises rapidly as animals initiate locomotion and begin navigation, then declines over several seconds. Such continuous ramping dynamics track elapsed time or traveled distance^26,27^. A manipulation that suppressed PyrUp activity also impaired task performance^26^, indicating that these dynamics contribute to behavior.

Here, we found that phasic LC activity at locomotion onset accompanied DA transients that were confined to micrometer-scale domains surrounding LC axons in CA1 and decayed over seconds. These localized DA transients, acting through D_1_-like receptors, preferentially amplified activity in nearby neurons exhibiting PyrUp dynamics. This amplification, in turn, biased the timing of reward-anticipatory behavior. Computational modeling indicated that this neuronal selectivity can arise from the coincidence of local DA exposure and strong ongoing activity in PyrUp neurons, without requiring dedicated targeting of a particular cell type. These findings reveal how a broadly projecting neuromodulatory system can selectively shape an ongoing computation: phasic LC activity determines when modulation occurs, spatially confined DA limits where it acts, and ongoing neuronal activity defines which nearby neurons are selectively modulated.

## Results

### LC run-onset activity reflects recent experience and shapes the timing of subsequent reward-anticipatory behavior

We trained head-fixed mice on a goal-directed navigation task, which we termed the virtual-reality integration (VRI) task (**Fig. 1A**)^26,27^. Each trial began with a brief visual start cue, after which the mouse navigated through a 180-cm landmark-free corridor. A reward was delivered only if the animal licked within an unmarked reward zone at the end of the corridor. Successful performance required the animal to integrate self-motion cues to estimate elapsed time or distance. Trained animals self-initiated running with variable latencies relative to the start cue, consistent with run onset marking the beginning of time/distance estimation^26^. When analysis was aligned to run onset, animals exhibited reward-anticipatory behavior: their running speed progressively slowed and licking increased as they approached the unmarked reward zone (**Fig. 1, B and C**). Because trained animals ran at relatively stereotyped speeds, elapsed time and traveled distance were strongly correlated^26^. Unless noted otherwise, we report results in the time domain, with key findings confirmed in the distance domain.

**Fig. 1.**
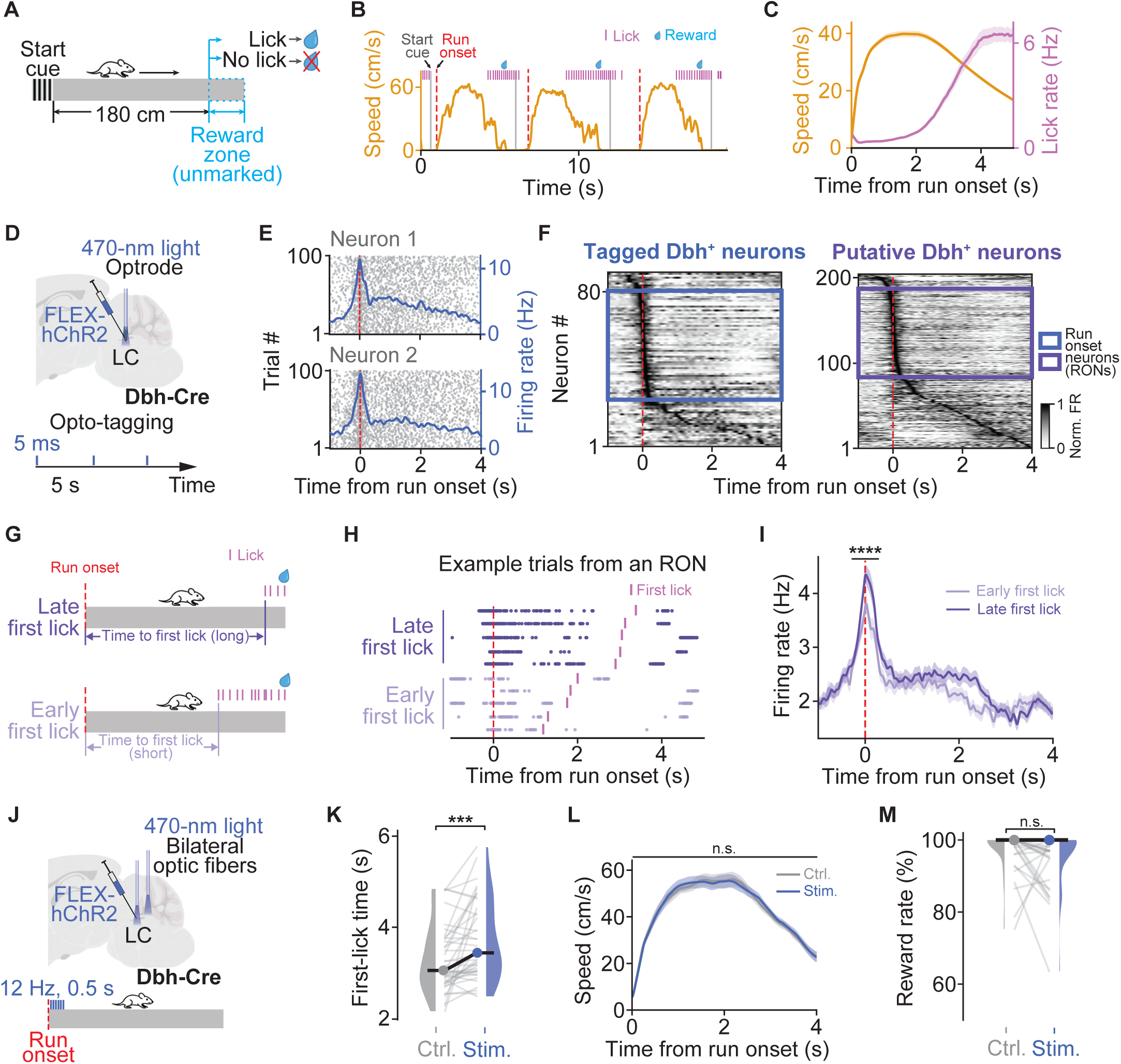
Phasic activity of LC Dbh^+^ neurons modulates action timing in a virtual-reality integration task. (**A**) Schematic of the virtual-reality integration task. (**B**) Three consecutive trials from a trained animal. Orange: running speed; gray: start cue; red: run onset; magenta: licks; blue: reward delivery. (**C**) Mean running speed (orange) and lick rate (magenta) across recordings. Shaded area represents SEM (lick-rate slope from 2–4 s after run onset: 3.03 ± 0.25 Hz/s; p = 4.30e-27; 25 animals, 127 recordings; one-sample *t*-test). (**D**) Schematic of extracellular recordings with opto- tagging of LC Dbh^+^ neurons. (**E**) Raster plots (gray) and trial-averaged firing rates (blue) aligned to run onset for two example opto-tagged Dbh^+^ neurons. Red dashed line, run onset. (**F**) Population heatmaps of trial-averaged firing rate aligned to run onset for tagged (blue, left) and putative (purple, right) Dbh^+^ neurons. Neurons are ordered by peak time. RONs are highlighted in colored squares. (**G**) Schematic defining first-lick time as the latency from run onset to the first anticipatory lick within a trial (early: first-lick time < 2.5 s; late: 2.5 s < first-lick time < 3.5 s). (**H**) Example trials illustrating early and late first-lick time. Raster shows spikes aligned to run onset (red dashed line) over multiple trials from an example RON; magenta marks indicate first-lick times. (**I**) Mean RON firing-rate profiles aligned to run onset for speed-matched early- versus late-first-lick trials (mean firing rate [-0.5, 0.5] s: early, median = 2.73 Hz, IQR = [1.85, 3.86] Hz; late, median = 3.02 Hz, IQR = [2.22, 4.16] Hz; p = 3.62e-05; Wilcoxon signed-rank test). n.s. p >= 0.05, * p < 0.05, ** p < 0.01, *** p < 0.001, **** p < 0.0001. (**J**) Schematic of bilateral optogenetic stimulation of LC Dbh^+^ neurons triggered at run onset (470 nm; 12 Hz for 0.5 s). (**K**) Summary across recordings of first-lick time for control versus stimulation trials. Gray lines connect paired control and stimulation trial-averages within a recording (control, median = 3.07 s, IQR = [2.81, 3.78] s; stimulation, median = 3.48 s, IQR = [3.06, 4.40] s; p = 2.58e-04; 12 animals, 54 recordings; Wilcoxon signed-rank test). (**L**) Mean running-speed profiles aligned to run onset for control and stimulation trials (control, median = 44.62 cm/s, IQR = [42.71, 46.48] cm/s; stimulation, median = 45.40 cm/s, IQR = [43.48, 46.10] cm/s; p = 0.71; Wilcoxon signed-rank test). (**M**) Reward rate across recordings (control, median = 100%, IQR = [99, 100]%; stimulation, median = 100%, IQR = [97, 100]%; p = 0.13; Wilcoxon signed-rank test).

We recorded LC activity extracellularly in Dbh-Cre mice, and identified dopamine-β- hydroxylase-positive (Dbh^+^) neurons by optogenetic tagging following ChR2 expression in the LC^28,29^ (**Fig. 1, D and E, and fig. S1, A and B**; see **Materials and methods**). We applied unsupervised clustering to spike auto-correlograms from all recorded neurons (**fig. S1C**; UMAP^30^; see **Materials and methods**). Untagged neurons clustered with the tagged population were identified as putative Dbh+ neurons. The waveforms and firing properties of both tagged and putative Dbh+ neurons resembled those previously reported for noradrenergic neurons^31–33^ (**fig. S1, B, D, and E**). The majority of Dbh^+^ neurons displayed a brief, phasic increase in activity at run onset (**Fig. 1, E and F**; tagged: 65.59%, 61 out of 93 neurons; putative: 53.85%, 112 out of 208 neurons; see **Materials and methods**). We term these “run-onset neurons” (RONs). In contrast, significantly fewer Dbh^+^ neurons showed responses aligned to the start cue or reward delivery (**fig. S1, F to H**; see **Materials and methods**).

Although LC activity has previously been linked to locomotion onset^16,17,34,35^, RON phasic responses do not simply reflect movement initiation. Response amplitude was not significantly correlated with running speed or acceleration around run onset (**fig. S2, A and B**; see **Materials and methods**). Moreover, when animals paused and spontaneously resumed running in the middle of a trial, the resulting spontaneous run onsets exhibited significantly smaller responses than trial-start run onsets, even after matching for running speed (**fig. S2C**; see **Materials and methods**). Thus, RON phasic responses were enhanced when locomotion onset marked the start of time/distance estimation.

RON phasic responses also depended on recent experience. They varied with the time elapsed between the previous reward and the next run onset, termed the reward-to-run interval. Longer intervals were correlated with higher baseline firing before run onset and larger phasic responses at the subsequent run onset (**fig. S3, C to F**; see **Materials and methods**). In a generalized linear model (GLM)^36^, reward-to-run interval remained a strong predictor of phasic response amplitude (**fig. S3, A and B**; see **Materials and methods**).

We next asked whether this trial-to-trial variation in RON response amplitude predicted when animals subsequently initiated reward-anticipatory licking (“first lick”). Responses were significantly larger on trials with later first licks than on trials with earlier first licks, and this difference persisted after matching the trial groups for running speed (**Fig. 1, G to I, and fig. S4A**; see **Materials and methods**). Conversely, RON responses did not differ significantly between trials with high and low running speed (**fig. S4B**). Together, these results indicate that RON response amplitude predicted subsequent action timing independently of running speed.

To test whether this phasic LC response can causally bias first-lick timing, we optogenetically stimulated Dbh^+^ neurons at trial-start run onset (12 Hz, 0.5 s; **Fig. 1J**). Compared to interleaved control trials, bilateral LC activation significantly increased both the time elapsed and distance traveled before the first lick (**Fig. 1K, and fig. S4C**), without significantly affecting running speed or reward rate (**Fig. 1, L and M**). Identical stimulation delivered at reward delivery did not reduce consummatory licking (**fig. S4D**), arguing against a non-specific suppression of licking. The same stimulation delivered mid- trial, at 120 cm, produced no detectable change in first-lick time, first-lick distance, or reward rate (**fig. S4E**). Thus, transient LC activation delayed reward-anticipatory action when delivered at trial-start run onset, but not later in the trial.

Together, these results suggest that the LC phasic response at trial-start run onset shapes the timing of reward-anticipatory action seconds later.

### LC phasic activity rapidly biases CA1 toward PyrUp dynamics

How can an LC phasic response lasting less than a second influence reward-anticipatory action several seconds later? CA1 ramping dynamics begin at run onset and evolve over the following seconds, providing a potential mechanism for extending the influence of this brief signal^26^. We therefore asked whether LC input shapes these ramping dynamics. We performed extracellular recordings in dorsal CA1 (**Fig. 2A**; see **Materials and methods**). Consistent with our previous work^26^, we identified two dominant functional pyramidal subpopulations based on their firing change before and after trial-start run onset. These subpopulations exhibited opposite ramping dynamics (**Fig. 2B**; see **Materials and methods**): PyrUp neurons (34.68%, 5065/14603; 57 animals, 201 recordings) exhibited a rapid synchronous increase in firing at run onset, followed by a gradual decay toward the unmarked reward zone; conversely, PyrDown neurons (18.32%, 2675/14603) showed an initial suppression followed by a gradual increase.

**Fig. 2.**
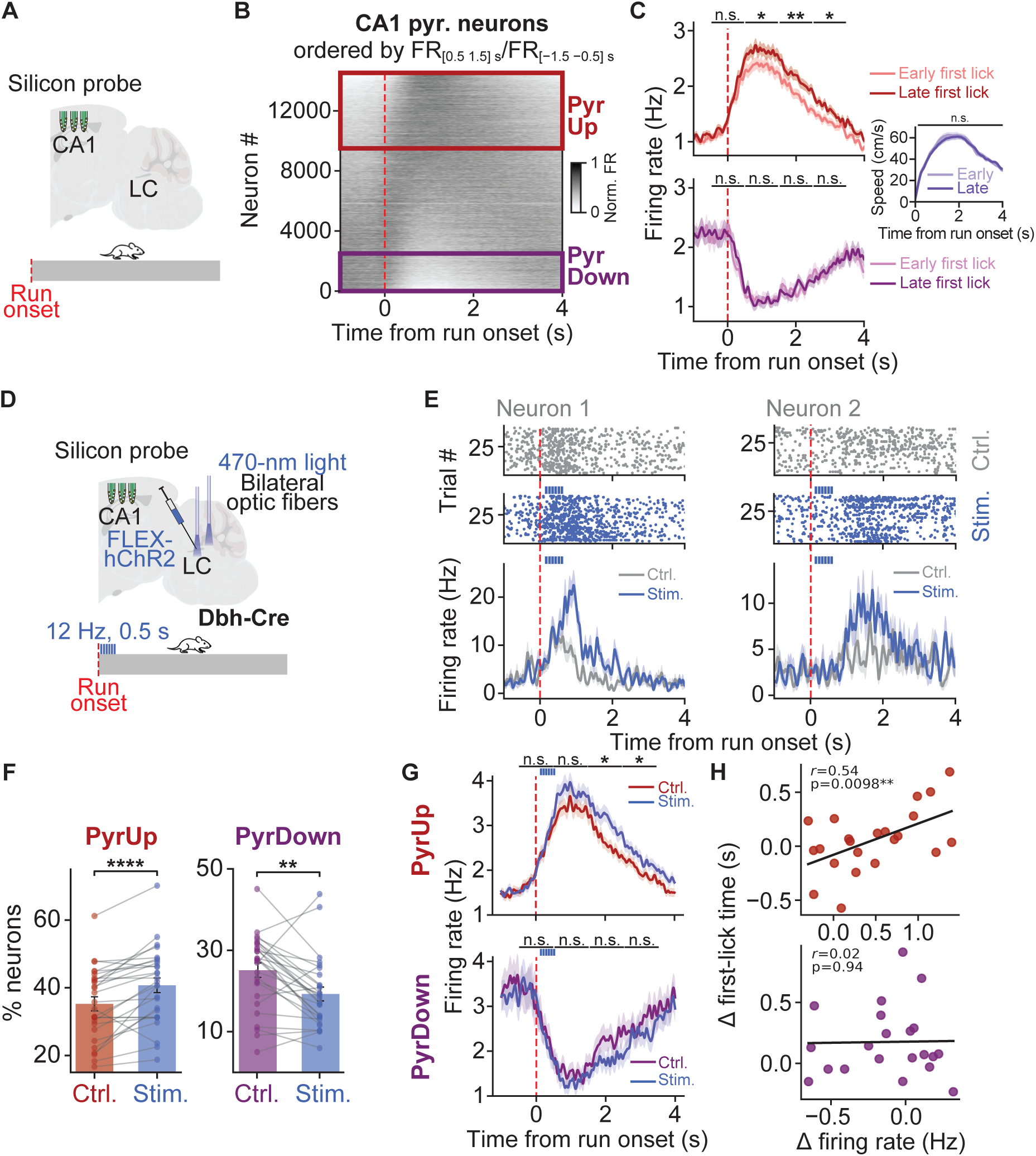
Phasic LC activation at trial-start run onset rapidly biases CA1 toward PyrUp dynamics. (**A**) Schematic of extracellular recordings in CA1 during the VRI task. (**B**) Population heatmap of CA1 pyramidal-neuron firing rates aligned to run onset. Neurons are ordered by their run-onset response index (R = FR_[0.5, 1.5] s_ / FR_[−1.5, −0.5] s_). PyrUp (red square): R>1.5; PyrDown (purple square): R<0.67. Red dashed line indicates run onset. (**C**) Mean firing-rate profiles for PyrUp (top) and PyrDown (bottom) neurons comparing early versus late first-lick trials after speed matching (17 animals, 25 recordings; independent *t*-test for each time bin); inset shows matched speed profiles (early, median = 43.85 cm/s, IQR = [41.78, 47.26] cm/s; late, median = 45.11 cm/s, IQR = [42.12, 46.92] cm/s; p = 0.15; Wilcoxon signed-rank test). (**D**) Schematic of unilateral CA1 recording combined with optogenetic activation of LC Dbh+ neurons triggered at trial-start run onset. (**E**) Two example CA1 pyramidal neurons showing rapid modulation during run-onset-triggered LC activation. Top, spike rasters over trials; bottom, mean firing rate profiles for stimulation (blue) and interleaved control trials (gray) (median activation latency = 0.96 s, IQR = [0.31, 2.00] s; 4 animals, 27 recordings). (**F**) Across recordings, percentages of PyrUp (left) and PyrDown neurons (right) in control and stimulation trials. Gray lines connect paired values within a recording (percentage: PyrUp, ctrl, median = 35.79%, IQR = [28.51, 42.64]%; LC stimulation, median = 41.86%, IQR = [32.29, 47.68]%; p = 2.55e-04; PyrDown, ctrl, median = 27.52%, IQR = [20.31, 31.02]%; LC stimulation, median = 18.09%, IQR = [13.29, 22.71]%; p = 2.05e-03; both Wilcoxon signed-rank tests). (**G**) Mean firing rate profiles for PyrUp (top) and PyrDown (bottom) neurons that maintained their classification in both control and stimulation trials (independent *t*- test for each time bin). (**H**) Scatter plots showing PyrUp (top) and PyrDown (bottom) stimulation- induced changes in firing rate (Δ firing rate in [0.5, 1.5] s) against changes in first-lick time (Δ first-lick time). Solid line indicates linear fit (ordinary least squares regression).

PyrUp neurons showed significantly higher firing rates on trials with late first licks than on trials with early first licks, after matching the trial groups for running speed. This firing-rate difference persisted for seconds after run onset (**Fig. 2C top and inset, and fig. S5A**; see **Materials and methods**). In contrast, PyrDown neurons did not differ significantly between these trial groups (**Fig. 2C bottom**).

Because both RON phasic response amplitude and PyrUp activity predicted first-lick timing, we next asked whether LC activity at trial-start run onset modulates PyrUp dynamics. We combined bilateral LC optogenetic activation with simultaneous silicon-probe recording in CA1 (**Fig. 2D**). At the single-neuron level, phasic LC activation modulated a subset of CA1 pyramidal neurons, with responses peaking approximately one second after run onset (peak latency: median = 0.96 s from run onset, IQR = [0.31, 2.00] s; **Fig. 2E**; protocol as in **Fig. 1J**). At the population level, LC activation significantly increased the percentage of PyrUp neurons while simultaneously decreasing the percentage of PyrDown neurons (**Fig. 2F, and fig. S5B**). To examine changes within these functional subpopulations, we analyzed neurons that maintained their PyrUp or PyrDown identity across control and stimulation trials. Among these neurons, LC activation significantly enhanced PyrUp responses over the ensuing seconds, while it did not significantly alter PyrDown responses (**Fig. 2G**). Across recording sessions, larger stimulation-induced increases in PyrUp responses (Δ firing rate in [0.5, 1.5] s) were associated with larger increases in first-lick time (**Fig. 2H, and fig. S5C**). Thus, run-onset LC activation preferentially enhanced PyrUp dynamics, and the magnitude of this enhancement tracked the subsequent delay in reward-anticipatory action.

Because somatic LC activation may indirectly influence CA1 through other downstream targets, we next asked whether direct LC–CA1 input can reproduce the CA1 neuronal effects. First, we examined the activity of LC–CA1 axons. Two-photon calcium imaging of axon-targeted GCaMP7b^37^ revealed that more than half of LC Dbh^+^ axon ROIs in CA1 exhibited calcium transients at trial-start run onset (363/669 ROIs, 54.26%; 3 animals, 50 recordings; **fig. S6, A to E**; see **Materials and methods**). Next, we optogenetically activated LC axons locally within CA1 at run onset. Unilateral axonal stimulation, with limited light power delivered to axon terminals (see **Materials and methods**), was unable to produce a detectable behavioral shift (first-lick time: ctrl., median = 2.75 s; stim., median = 2.91 s; p = 0.23; Wilcoxon signed-rank test; 5 animals, 29 sessions). Nevertheless, it recapitulated the key neuronal effects observed with LC somatic stimulation: it rapidly modulated a subset of CA1 pyramidal neurons and increased the percentage of PyrUp neurons (**fig. S5, D to H**).

Together, these findings support a model in which the LC–CA1 projection extends the influence of a phasic run-onset signal over the following seconds by promoting PyrUp dynamics, thereby delaying reward-anticipatory action. This raises the question of which transmitter mediates this seconds-timescale LC–CA1 effect.

### LC phasic activity delivers spatially confined dopamine transients to CA1

Although the LC is canonically noradrenergic, its hippocampal terminals may also release dopamine^14,15,18,19^. Yet the spatial organization of these signals remains unknown. To test this directly, we expressed the genetically encoded DA sensor dLight3.6^38,39^ in CA1 and tdTomato-ChrimsonR in LC Dbh^+^ neurons. We then optogenetically stimulated LC while imaging CA1 with two-photon microscopy in awake, head-fixed mice (**Fig. 3A**). Phasic LC activation produced a rapid, robust increase in the field-averaged dLight signal. No corresponding transient occurred in the tdTomato channel labeling LC axons, indicating that the dLight signal was not caused by motion artifacts (**Fig. 3, B and C**; see **Materials and methods**). Because dLight retains some sensitivity to NE^38,39^, we next depleted NE by inhibiting its synthesis from DA with nepicastat^40^ (i.p.; see **Materials and methods**). Phasic LC activation continued to evoke robust dLight transients following nepicastat treatment (**fig. S7, A and B**), consistent with DA as a major contributor to the LC-evoked signals.

**Fig. 3.**
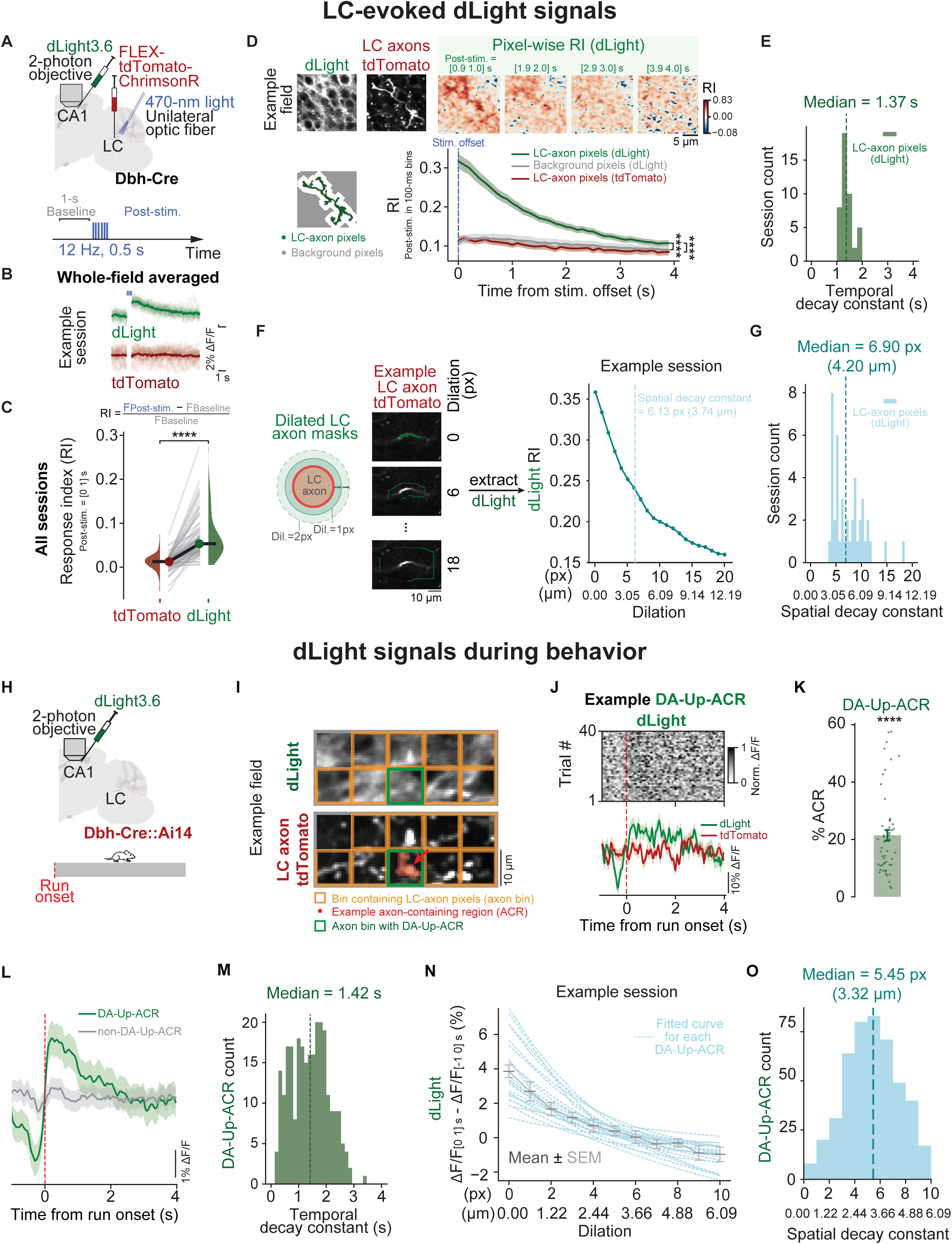
Rapid, spatially localized dLight signals around LC axons in CA1. (**A**) Experimental schematic for two-photon imaging of dLight3.6 in CA1 during optogenetic activation of LC Dbh^+^ neurons while animals were head-fixed. Stimulation at 470 nm was used, which reduces opsin sensitivity but minimizes PMT interference. (**B**) Example recording showing whole-field-averaged signals from dLight (green) and reference channel (tdTomato, red) aligned to stimulation (blue bars) (baseline vs. post-stimulation [0, 1] s: dLight, p = 3.81e-06; tdTomato control, p = 0.40; 19 stimulation epochs; Wilcoxon signed-rank test). (**C**) Summary across recordings of tdTomato (control) and dLight response indices in CA1. Top: formula for calculating response index (RI) (tdTomato RI, median = 0.01, IQR = [0.00, 0.02]; dLight RI, median = 0.05, IQR = [0.04, 0.08]; p = 3.75e-14; 6 animals, 77 recordings; Wilcoxon signed-rank test). (**D**) Spatial distribution of stimulation-evoked dLight signals relative to LC axons, analyzed at the pixel level across the field of view. Top, example subfield containing an LC axon, showing the dLight channel and the tdTomato channel (left two panels); pixel-wise RI maps at the indicated post-stimulation times relative to stimulation offset are shown for the same region (right). Bottom left: classification of LC-axon pixels (dark green) vs background pixels (gray). Bottom right: RI traces are aligned to stimulation offset and averaged across recordings, showing dLight signals extracted from LC-axon pixels (green) versus background pixels (gray) across the whole field, together with tdTomato signals extracted from LC-axon pixels (red). (**E**) Histogram of temporal decay constants obtained by fitting an exponential to the post-stimulation RI time course measured in LC-axon pixels for each recording (median decay constant=1.37 s). (**F**) Left, schematic illustrating the spatial dilation analysis: concentric 1-pixel rings are defined at increasing distances from the LC-axon mask to measure dLight signal as a function of distance. An example LC axon with its corresponding dilated masks is shown. Right, example session, mean dLight RI within each successive 1-pixel ring as a function of distance from LC axon pixels, quantified using the [0, 1] s window after stimulation offset as the post-stim window. (**G**) Histogram of spatial decay constants obtained by exponential fits to the RI-versus-dilation-distance profiles for each recording. (**H**) Experimental schematic for two-photon imaging of dLight with tdTomato-labeled LC axons in CA1 during the VRI task. (**I**) Example showing how dLight signal was extracted from LC axon segments. The imaging field was divided into approximately 10×10 µm bins (dLight channel, top; tdTomato channel, bottom). Bins containing tdTomato-positive LC axon area (“axon bin”) are outlined in orange. Within each axon bin, the tdTomato-positive area that also expresses dLight defines the axon-containing region (ACR). One axon bin that contains an ACR with a significant run-onset dLight transient (“DA-Up-ACR”) is outlined in green; the ACR itself is indicated by a red arrow. (**J**) dLight signal from an example DA-Up-ACR, from the bin outlined in the green square in (**I**). Top, trial-by-trial dLight signal aligned to run onset. Bottom, dLight and tdTomato traces averaged across trials. (**K**) Percentage of ACRs classified as DA-Up-ACRs. Each dot represents one recording (percentage of DA-Up-ACRs, 21.42 ± 1.86%; p = 3.50e-12; 4 animals, 57 recordings; one-sample *t*-test against 5%). (**L**) dLight trace averaged across all DA-Up-ACRs (green) and non- DA-Up-ACRs (gray). (**M**) Histogram of temporal decay constants for dLight signals from DA- Up**-**ACRs (median = 1.42 s). (**N**) Example session showing the change in dLight signal within each dilated DA-Up**-**ACR mask, computed as post-run-onset minus pre-run-onset ΔF/F and plotted as a function of dilation distance from the original DA-Up-ACR. Blue traces show the fitted exponential decay for individual DA-Up-ACRs. Gray points and error bars indicate the mean ± SEM across all DA-Up-ACRs at each dilation distance. (**O**) Histogram of spatial decay constants obtained by exponential fits in (**N**) for each DA-Up-ACR (median decay constant = 3.32 μm).

We next asked whether LC-evoked dLight signals diffuse broadly through extracellular space^5–7^, or remain confined near axonal terminals. dLight signals were strongest at LC axons and decayed quickly with distance (**Fig. 3D top**; see **Materials and methods**), with a median spatial decay constant of 4.20 μm during the first second after stimulation (**Fig. 3, F and G**; see **Materials and methods**). Transients were detected across the majority of LC axonal ROIs (median = 87.5% of axonal ROIs per recording; **fig. S7C**; see **Materials and methods**), and their magnitude did not correlate with dLight expression levels (**fig. S7D**; see **Materials and methods**). These transients were also brief, decaying with a median time constant of 1.37 s at LC axons (**Fig. 3, D bottom and E**; see **Materials and methods**). Together, these results indicate that phasic LC activation generates DA signals that remain confined to micrometer-scale domains surrounding the axons and persist for seconds. These seconds-long signals overlap with the period during which LC activity shapes PyrUp dynamics and action timing.

We next tested whether dopaminergic signaling is necessary for task performance. Bilateral infusion of the D_1_-like dopamine receptor antagonist SCH-23390 in CA1^41–43^ significantly reduced running speed, increased licking shortly after run onset, and decreased reward rate (**fig. S8, A and B bottom**). In contrast, bilateral infusion of the α₁-adrenergic antagonist prazosin or the β-adrenergic antagonist propranolol^44–49^ failed to significantly affect running speed, licking patterns, or reward rate (**fig. S8, A and B top and middle**). These distinct behavioral effects collectively implicate D_1_-like signaling in CA1 as critical for task performance.

### Spatially localized dopamine transients occur at run onset during behavior

Having established that optogenetically evoked LC activity generates spatially confined dLight transients around LC–CA1 axons, we next asked whether comparable signals occur during task performance. We simultaneously imaged dLight signals and tdTomato-labeled LC axons in CA1 of Dbh-Cre::Ai14 mice (**Fig. 3H, and fig. S9A**). To map dLight signals along LC axons, we divided each imaging field into small spatial bins (∼10×10 μm) and defined the tdTomato-positive axonal area within each bin as an axon-containing region (ACR; **Fig. 3I**; see **Materials and methods**).

A subset of ACRs exhibited significant dLight transients at trial-start run onset, which we termed DA-Up-ACRs (**Fig. 3, J and L;** see **Materials and methods** for motion correction using the tdTomato channel). These transients peaked after LC–CA1 axonal calcium activity (peak time: LC–CA1 run-onset-active axons, median = 0.37 s, IQR = [0.27, 0.86] s; dLight, median = 0.93 s, IQR = [0.34, 1.80] s; **fig. S9C**), consistent with catecholamine release following LC axonal activation.

These run-onset dLight transients were spatially confined to micrometer-scale domains surrounding DA-Up-ACRs, with a median spatial decay constant of 3.32 μm during the first second after run onset (**Fig. 3, N and O, and fig. S9B**; see **Materials and methods**). The transients decayed with a median time constant of 1.42 s (**Fig. 3M**; see **Materials and methods**).

We next examined how run-onset dLight signals were distributed among LC–CA1 axonal segments. DA-Up-ACRs comprised only 21.42 ± 1.86% of all ACRs (**Fig. 3K**), with the percentage exceeding chance levels in 48/57 recordings (see **Materials and methods**). Across the imaging field, DA-Up-ACRs were no more clustered than randomly selected ACRs (**fig. S9D**). Thus, run-onset dLight signals were not only spatially confined near axons but also restricted to discrete axonal segments. This variation among axonal segments was unlikely to reflect variation in dLight expression, as ACR response magnitude did not correlate with expression level (**fig. S9E**). Compared with optogenetically evoked signals, run-onset dLight signals were detected at fewer axonal segments.

In summary, dLight transients arose at discrete LC–CA1 axonal segments at run onset, when animals began time/distance estimation in the VRI task. These transients remained confined to micrometer-scale domains and lasted seconds. Together with the optogenetically evoked dLight transients, these signals suggest that LC phasic activity produces a catecholamine transient in CA1, with DA as a major contributor.

### Spatially localized dopamine preferentially promotes PyrUp dynamics through D_1_- like receptors

Because LC-derived dLight transients were confined to micrometer-scale domains surrounding discrete LC–CA1 axonal segments, we reasoned that only neurons near DA- Up-ACRs would be exposed to these transients. To identify these neurons and examine their activity, we co-expressed dLight3.6 and the red-shifted calcium indicator jRGECO1a^50^ in CA1 and performed dual-color two-photon imaging (**Fig. 4A**). Because LC axons were not labeled in this preparation, we used the dLight signal measured at the perisomatic membrane to assess local DA exposure at each soma (see **Materials and methods**).

**Fig. 4.**
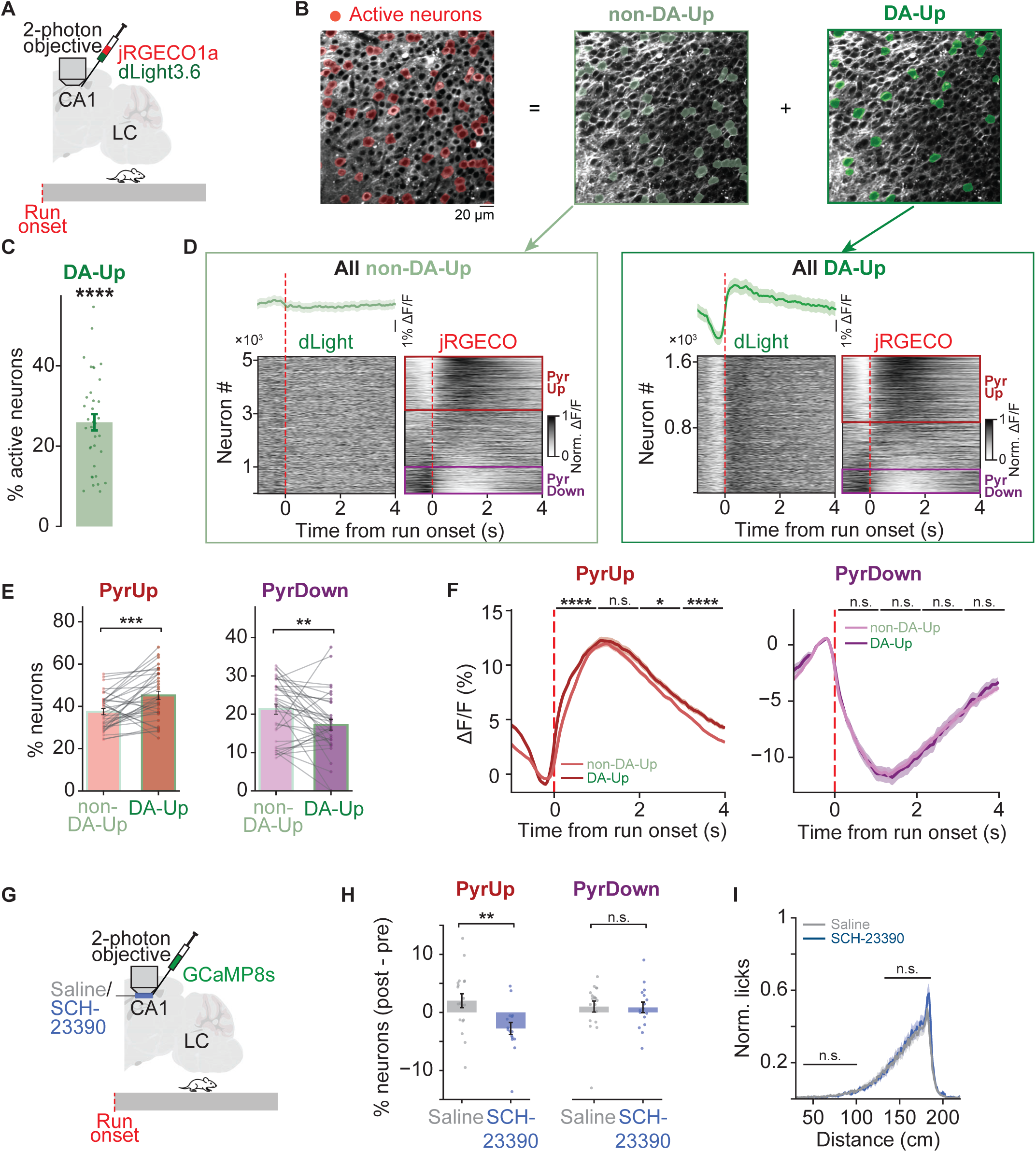
Perisomatic dLight transients are preferentially associated with recruitment and amplification of PyrUp dynamics. (**A**) Schematic of dual-color two-photon imaging of dLight and jRGECO in CA1 during the VRI task. (**B**) Example field of view. Left: jRGECO image with active neurons highlighted in red. Middle/right: the same field in the dLight channel with active neurons separated into non-DA-Up (light green, middle) and DA-Up (dark green, right) groups. (**C**) Percentage of active neurons classified as DA-Up. Each dot represents one recording (25.94 ± 2.02%; p = 4.85e-12; 5 animals, 35 recordings; one-sample *t*-test against 5%). (**D**) For all non-DA-Up neurons (left group, light green box) and all DA-Up neurons (right group, dark green box), paired heatmaps show dLight signal (left column) and jRGECO calcium signal (right column) aligned to run onset. Neurons are ordered by their run-onset calcium responses. Red and purple squares indicate neurons classified as PyrUp and PyrDown, respectively. dLight traces averaged for each group are shown above the corresponding heatmaps. (**E**) Percentages of PyrUp (left) and PyrDown (right) neurons in DA-Up versus non-DA-Up groups, summarized across recordings. Gray lines connect paired values within a recording (percentage of PyrUp: non-DA-Up, 37.46 ± 1.43%; DA-Up, 45.17 ± 1.97%; p = 1.92e- 04; percentage of PyrDown: non-DA-Up, 21.33 ± 1.31%; DA-Up, 17.23 ± 1.38%; p = 8.23e-03; both Wilcoxon signed-rank tests). (**F**) Calcium traces averaged across PyrUp (left) and PyrDown (right) neurons, split by DA-Up (darker traces) and non-DA-Up (lighter traces) groups. ΔF/F was baseline-corrected by subtracting the mean signal in the [−0.5, 0] s window before run onset. Asterisks indicate time bins with significant differences between groups (independent *t*-test). (**G**) Schematic of local pharmacology during CA1 two-photon imaging: saline or SCH-23390 was infused through the imaging window while recording CA1 activity. (**H**) Change in the percentage of PyrUp (left) and PyrDown (right) neurons following saline (gray) or SCH-23390 (dark blue) infusion, relative to pre-infusion sessions (PyrUp change: saline, 2.03 ± 1.19%; SCH-23390, -2.76 ± 1.03%; p = 4.53e-03; PyrDown change: saline, 1.01 ± 0.96%; SCH-23390, 0.87 ± 0.92%; p = 0.92; 8 animals, 19 recordings for saline, 16 recordings for SCH-23390; all independent t-tests). (**I**) Licking profiles for saline versus SCH-23390 infusion sessions (lick rate was normalized to the maximum lick rate across both conditions: licks [30, 100] cm: saline, median = 0.011, IQR = [0.004, 0.021]; drug, median = 0.011, IQR = [0.005, 0.018]; p = 0.82; anticipatory licking [120, 180] cm: saline, median = 0.210, IQR = [0.173, 0.319]; drug, median = 0.229, IQR = [0.201, 0.278]; p = 0.79; all Wilcoxon rank-sum tests).

We identified “active neurons”, defined as pyramidal neurons with clear somatic calcium activity (**Fig. 4B**; see **Materials and methods**). For each active neuron, we then extracted a motion-corrected perisomatic dLight trace and classified the neurons as “DA-Up” if the dLight trace showed a significant increase at run onset (see **Materials and methods**). Of active neurons, 25.94 ± 2.02% were DA-Up (**Fig. 4C**), with the percentage significantly exceeding chance levels in 35/35 recordings (see **Materials and methods**).

DA-Up neurons were significantly more likely to exhibit PyrUp dynamics and less likely to exhibit PyrDown dynamics, compared to the remaining active neurons (“non-DA-Up”; **Fig. 4, D and E**; see **Materials and methods**). DA-Up neurons exhibiting PyrUp dynamics also showed significantly larger calcium responses after run onset than non-DA-Up PyrUp neurons (**Fig. 4F left**). In contrast, calcium responses did not differ between DA-Up and non-DA-Up neurons exhibiting PyrDown dynamics (**Fig. 4F right**). Together, perisomatic dLight transients at run onset were associated with both preferential recruitment into PyrUp dynamics and amplification of PyrUp response magnitude. This pattern was consistent with our optogenetic finding that phasic LC activation increased both the prevalence and amplitude of PyrUp dynamics (**Fig. 2, F and G**).

Finally, we tested whether D_1_-like receptor signaling promotes PyrUp dynamics. We imaged CA1 activity with GCaMP8s^51^ while locally infusing SCH-23390 through the imaging window^52^ (**Fig. 4G**). To minimize effects on neural activity arising from behavioral changes, we used a lower antagonist dose that did not significantly alter running speed, licking patterns or reward rate (**Fig. 4I, and fig. S10A**). At this dose, SCH-23390 significantly reduced the percentage of PyrUp neurons (**Fig. 4H**). In contrast, blocking α_1_- or β-adrenergic receptors produced no significant change in PyrUp/PyrDown prevalence, even at the high dose used in the behavioral experiments (**fig. S10, A and B**). Together, our findings support a model in which spatially localized LC-derived DA signals preferentially promote PyrUp dynamics through D_1_-like receptors (**Fig. 5D**).

**Fig. 5.**
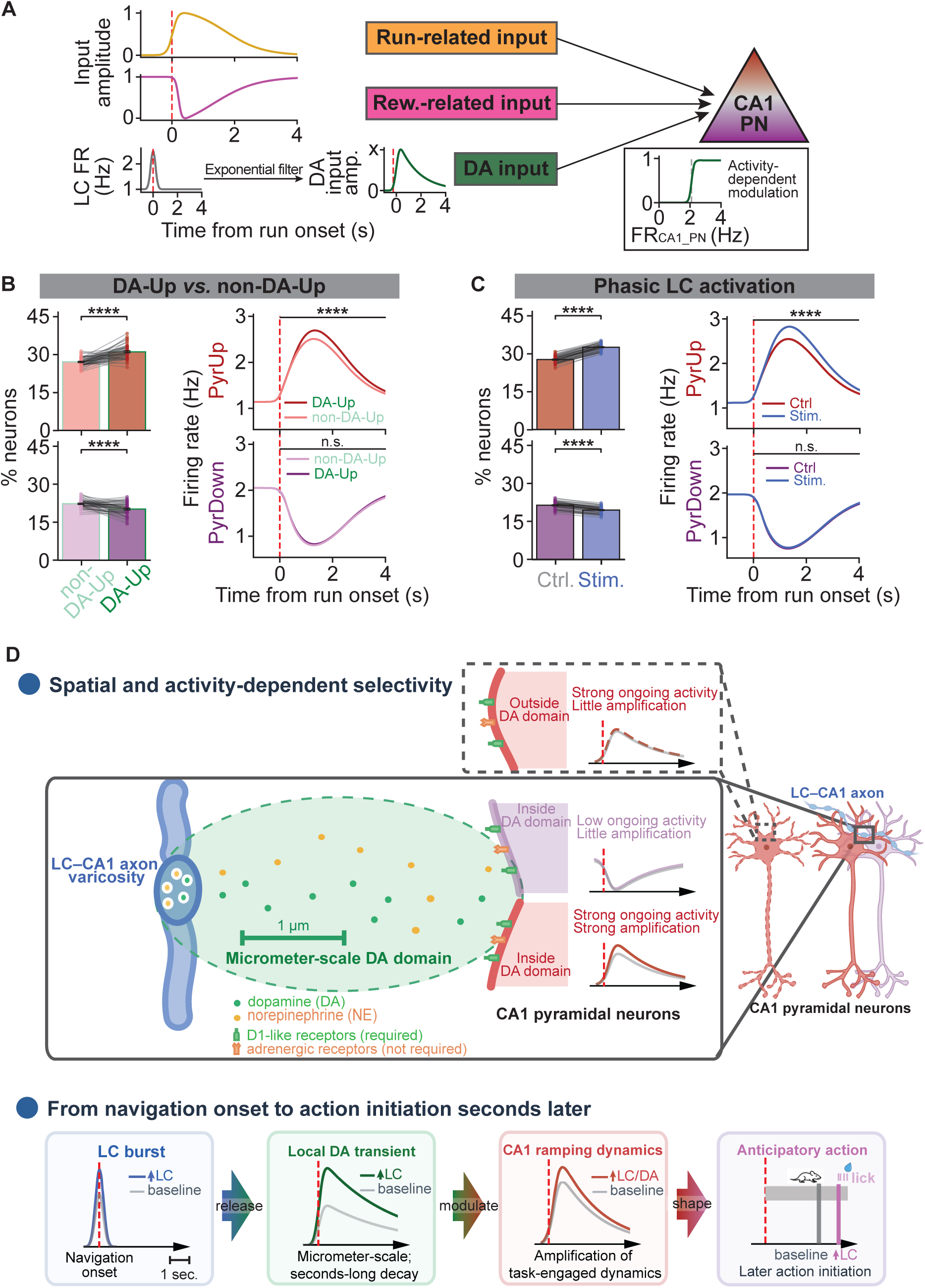
An LC–dopamine–CA1 model reproduces selective PyrUp enhancement. (**A**) Schematic of an LC–dopamine–CA1 model. Each CA1 pyramidal neuron receives three inputs (run-related, reward-related, and DA inputs). The DA input modulates a subset of neurons in an activity-dependent manner. (**B**) Left: percentage of PyrUp (top) and PyrDown (bottom) neurons among DA-Up (DA receiving) versus non-DA-Up model neurons. Gray lines connect paired results in individual simulation runs (percentage: PyrUp, non-DA-Up, median=27.00%, IQR=[26.00, 28.25]%; DA-Up, median=30.83%, IQR=[29.33, 32.67]%; p=9.53e-09; PyrDown, non-DA-Up, median=22.36%, IQR=[21.29, 23.25]%; DA-Up, median=20.33%, IQR=[18.67, 22.00]%; p=2.63e-05; both Wilcoxon signed-rank tests). Right: mean firing-rate profiles for PyrUp (top) and PyrDown (bottom) neurons in DA-Up versus non-DA-Up groups (firing rate [0, 4] s: PyrUp, non-DA-Up, median=1.96 Hz, IQR=[1.91, 2.00] Hz; DA-Up, median=2.08 Hz, IQR=[2.03, 2.13] Hz; p=2.44e-11; PyrDown: non-DA-Up, median=1.29 Hz, IQR=[1.24, 1.34] Hz; DA-Up, median=1.30 Hz, IQR=[1.26, 1.34] Hz; p=0.39; both Mann-Whitney U tests). (**C**) Same as (**B**) but comparing phasic LC activation trials versus control trials (percentage: PyrUp, ctrl, median=28.30%, IQR=[27.20, 29.32]%; LC stimulation, median=33.45%, IQR=[32.17, 34.30]%; p=7.53e-10; PyrDown, ctrl, median=21.80%, IQR=[20.92, 22.60]%; LC stimulation, median=19.90%, IQR=[18.70, 20.60]%; p=7.43e-10; both Wilcoxon signed-rank tests. Firing rate [0, 4] s: PyrUp, ctrl, median=1.97 Hz, IQR=[1.93, 2.02] Hz; LC stimulation, median=2.15 Hz, IQR=[2.11, 2.19] Hz; p=5.64e-17; PyrDown: ctrl, median=1.21 Hz, IQR=[1.17, 1.25] Hz; LC stimulation, median=1.23 Hz, IQR=[1.18, 1.27] Hz; p=0.15; both Mann-Whitney U tests). (**D**) A conceptual model illustrating micrometer-scale LC dopamine selectively amplifies active CA1 neurons and shapes action timing.

### A model of LC-derived dopamine signaling reproduces selective PyrUp enhancement

An unresolved question is why among neurons exposed to the local LC-derived DA, PyrUp neurons are preferentially enhanced. Studies of D_1_-like receptor physiology suggest one possibility: DA may amplify neurons that are already firing strongly when the transient arrives^53,54^.

DA transients and PyrUp enhancement both occurred within seconds of run onset. To test whether activity-dependent dopamine modulation can account for the observed selectivity on this timescale, we constructed a minimal phenomenological network model. CA1 neurons received run-related and reward-related excitatory inputs with random weights, from which PyrUp and PyrDown dynamics emerged. To represent spatially restricted dopamine exposure, phasic LC activity at run onset triggered a DA transient targeting a randomly selected subset of neurons, independently of their PyrUp or PyrDown identity. The transient began to modulate neuronal firing after a brief delay, so that its effect occurred within the seconds-long window observed experimentally. The strength of the modulation increased with each neuron’s ongoing firing rate (**Fig. 5A**; see **Materials and methods**).

The model reproduced the key experimental findings across three conditions. First, among PyrUp neurons, those exposed to DA showed greater firing rate enhancement than those that were not exposed; among PyrDown neurons, DA exposure produced no clear difference. DA-exposed neurons were also more likely to exhibit PyrUp dynamics and less likely to exhibit PyrDown dynamics (**Fig. 5B**). This result reproduced the differences between DA-Up and non-DA-Up neurons observed experimentally (**Fig. 4, E and F**). Second, simulated optogenetic LC activation at run onset increased PyrUp prevalence and enhanced firing among neurons that maintained their PyrUp identity across stimulation and control trials (**Fig. 5C**). This is consistent with the optogenetic results in **Fig. 2, F and G**. Third, simulating D_1_-like receptor blockade by removing the dopamine drive reduced PyrUp prevalence (**fig. S11**), consistent with the SCH-23390 results shown in **Fig. 4H**.

Together, this model demonstrated that activity-dependent dopaminergic modulation selectively promotes PyrUp dynamics (**Fig. 5D**). This selectivity requires no specific anatomical targeting of PyrUp neurons by LC axons; it emerges naturally from the coincidence of local DA exposure and strong ongoing neuronal activity in PyrUp but not PyrDown neurons. In this account, spatial confinement limits which neurons encounter dopamine, whereas ongoing activity determines which of those neurons are strongly modulated. Together, these properties allow a brief LC signal to shape CA1 dynamics over the seconds preceding reward-anticipatory action.

## Discussion

Neuromodulatory systems project widely across the brain, and the transmitters they release are thought to spread through surrounding tissue to influence many neurons within a target region^5–7^. Yet our findings reveal that the canonically noradrenergic LC can signal with unexpected spatial and functional selectivity: LC phasic activity produces dopamine transients confined to micrometer-scale domains around LC axons, preferentially amplifies activity in CA1 neurons engaged in goal-directed navigation, and shapes when animals initiate action seconds later. Modeling shows that this selectivity can emerge without dedicated wiring: among neurons exposed to a localized dopamine signal, those with strong ongoing activity are preferentially amplified.

The run-onset LC response was not simply a generic locomotor signal. It was stronger when run onset marked the beginning of time/distance estimation, varied with recent trial history, and predicted when animals subsequently initiated reward-anticipatory action. Briefly enhancing this response delayed action initiation. Together with the effects of LC activation on CA1 PyrUp dynamics, these findings suggest that phasic LC activity influences how task-engaged dynamics in the downstream circuits unfold over the following seconds. This extends the established functions of LC–CA1 signaling from memory-related processes spanning minutes to hours^14,15,17,55^ to the regulation of neuronal dynamics during ongoing behavior.

Several observations support the LC as a major source of the run-onset dopamine signal. Brief optogenetic LC activation evoked dLight transients in CA1 that persisted after inhibition of NE synthesis. Although incomplete inhibition and residual dLight sensitivity to NE leave some ambiguity, NE alone cannot readily account for the evoked signal. During behavior, run-onset dLight transients arose around LC axons and peaked shortly after LC axonal calcium activity. Moreover, VTA projections to dorsal CA1 are sparse relative to LC input^14–16,56^, and VTA axon activity tends to ramp toward reward^16,57–61^. These observations favor an LC origin at run onset but do not exclude VTA contributions at other stages of the task. Receptor pharmacology further supports a functional role for dopamine: blocking D_1_-like, but not α_1_- or β-adrenergic, receptors disrupted task performance, and a lower dose of the D_1_-like antagonist that did not significantly alter measured behavior still reduced PyrUp prevalence.

The spatial profile of the signal provides one source of specificity. The dLight transients declined sharply within a few micrometers of LC axons, plausibly reflecting local uptake and clearance^62–64^ and physical limits on extracellular diffusion^65,66^. They were also detected only around discrete axonal segments, consistent with spatial compartmentalization reported for VTA axons in cortex^38^ and striatum^67^.

Localized dLight transients preferentially accompanied enhanced activity in neurons with PyrUp dynamics. A potential explanation for this selectivity comes from evidence that D_1_- like signaling preferentially amplifies neurons that are already depolarized or firing^53,54^. Our model reproduced the observed selectivity without targeted innervation of PyrUp neurons, consistent with PyrUp and PyrDown populations being defined by their activity patterns rather than known molecular identity^26^.

How enhanced PyrUp activity contributes to action timing remains unresolved. Greater PyrUp activity predicted later initiation of reward-anticipatory licking, and LC activation both enhanced PyrUp dynamics and delayed action. These aligned effects motivate the hypothesis that modulation of PyrUp activity contributes to the behavioral change. Potential downstream readouts may include thresholding declining PyrUp activity, integrating it over time, or comparing the dynamics of PyrUp and PyrDown populations. These mechanisms differ in their downstream circuit requirements and remain to be tested.

More broadly, these findings show that broad anatomical reach does not preclude specificity. LC-derived dopamine operates over seconds and micrometers, a signaling regime intermediate between fast synaptic transmission and slower, spatially diffuse neuromodulation. Within this regime, selectivity can emerge from where a transmitter is available and which nearby neurons are active, rather than being fully specified by dedicated wiring. This organization could allow the same broadly projecting neuromodulatory system to influence different neuronal ensembles at different moments as local circuit activity changes.

## Materials and methods

### Experimental protocols and subjects

#### Animals

This study used both male and female mice (age > 7 weeks). Four mouse lines were used: C57BL/6J (JAX #000664), Dbh-Cre (JAX #033951)^68^, Drd1-Cre (MMRRC #030989), and Ai14 (JAX #007914)^69^. All procedures were performed in accordance with protocols approved by the Institutional Animal Care and Use Committee at Max Planck Florida Institute for Neuroscience. Mice were housed on a 12:12 reverse light-dark cycle, and behavior was tested during the dark phase.

#### Viral vectors

For optogenetic experiments, we used AAV5-Ef1a-DIO-ChR2(H134R)-mCherry (Addgene, titer 1.4e+13 vg/mL and 2.5e+13 vg/mL) and AAV5-Syn-FLEX-ChrimsonR-tdTomato (Addgene, titer 2.4e+13 vg/mL). For two-photon imaging, we used AAV5- CAG-DIO-axon-jGCaMP7b-WPRE-hGH-polyA (BrainVTA, titer 1.1e+13 vg/mL), AAV1-ihsyn-tTa-sv40/TRE-dLight3.6-minWPRE (gifts from Lin Tian’s Lab; titer 3.0e+11 vg/mL), AAV1-Syn-NES-jRGECO1a-WPRE-SV40 (Addgene; titer 2.2e+13 vg/mL), and AAV9-syn-jGCaMP8s-WPRE (Addgene, titer 2.4e+13 vg/mL).

#### Stereotaxic surgeries

Mice were anesthetized with isoflurane (3% induction; 1–2% maintenance) and mounted in a stereotaxic frame, with body temperature maintained at ∼37 °C using a heated pad. Eye ointment (Soothe, Bausch+Lomb, Laval, Canada) was applied to prevent corneal drying.

All mice were implanted with a headplate for head fixation. Electrophysiology, optogenetics, and pharmacology experiments used a lightweight plastic headplate; two-photon imaging experiments used a metal headplate. Headplates were secured to the skull with dental cement after standard skull preparation to maximize stability.

#### Viral injections

Viruses were delivered through pulled glass micropipettes using stereotaxic injections at the following target coordinates: LC: AP −5.45, ML ±1.20, DV −3.00 mm from the dura; dorsal hippocampus (CA1 region): AP −1.70, ML ±1.40, DV −2.10 mm from the dura. Pipettes were lowered slowly to the target depth to minimize tissue damage and backflow. Injections were performed at low flow rates. Typical volumes were 250 nL (LC) and 50 nL (hippocampus) per hemisphere.

#### Craniotomies and implantation

For electrophysiology, craniotomies were made over the LC (AP −5.45, ML ±1.00 mm) or dorsal hippocampus (AP −1.70, ML ±1.40 mm). For optogenetics, optic fibers were implanted to target the LC (AP −5.45, ML ±1.35, DV −2.30 mm from the dura), with a 10° tilt toward the midline. For optogenetics accompanying two-photon imaging, see **Two-photon imaging window**. For pharmacology, bilateral guide cannulae were implanted over the dorsal hippocampus at the same coordinates used for hippocampal viral injections. For optogenetic and pharmacological implants, a light-blocking “crown” (copper-foil shield painted black) was built around the implant to protect the optic fibers or the cannulae.

#### Two-photon imaging window

For two-photon imaging, a chronic imaging window (a cannula ∼3 mm in diameter, ∼1.5 mm in depth, on one end glued with a ∼3 mm diameter glass window) was implanted over the right dorsal hippocampus. After opening a circular craniotomy, the cortex and most of the fiber layers above dorsal CA1 were carefully removed, and the window was lowered to the CA1 surface and secured with C&B Metabond Adhesive Luting Cement (Parkell, New York, USA)^70^ .

For two-photon experiments combining imaging with LC optogenetics, an optic fiber was implanted to target the ipsilateral LC with a rostral tilt of 25° to avoid interference with the objective (AP −6.66, ML −1.0, DV −2.67 mm from the dura).

#### Post-operative analgesia

Post-operative analgesia included long-acting buprenorphine (SR or ER formulation, 1.0 mg/kg) and sustained-release meloxicam (0.5 mg/kg), administered subcutaneously.

#### Histology

Mice were deeply anesthetized with isoflurane and transcardially perfused with 20 mL of 1× PBS followed by 20 mL of 4% paraformaldehyde (PFA). Brains were extracted and post-fixed in 4% PFA for 24 h at 4 °C, then transferred to 1× PBS for storage until sectioning.

Fixed brains were embedded in 3% agarose and sectioned on a vibratome into 50-μm coronal slices. Sections were either mounted directly onto slides or collected in 24-well plates for immunohistochemistry.

To verify viral targeting in the LC, free-floating sections were processed for immunohistochemistry. Sections were permeabilized and blocked in 1× PBS containing 1% Triton X-100 and donkey serum, then incubated overnight at 4 °C with an anti–tyrosine hydroxylase (TH) primary antibody (AB152, MilliporeSigma; 1:500). After washes in 1× PBS, sections were incubated for 1 h with an Alexa Fluor 488–conjugated secondary antibody (Jackson ImmunoResearch; 1:1000), washed again, mounted onto slides, and counterstained with DAPI. Images were acquired on a Zeiss confocal microscope (LSM 980).

#### Behavior

The behavioral apparatus consisted of a projector or screens and a head-fixation stage mounted with a treadmill. Infrared illumination and cameras were used for behavioral tracking. Treadmill rotation was recorded with an encoder and converted into distance for analysis. The virtual environment was controlled by a Teensy 3.5 microcontroller on a custom Arduino board using in-house scripts, and all behavioral variables were synchronized through the microcontroller.

Animals were placed on water restriction at least three days after surgery. Daily water intake was maintained at 0.6–1.0 mL, with temporary adjustments as needed to maintain health. On the first day of restriction, animals were handled and habituated to the head-fixed setup. Animals were then trained to run while head-fixed on the treadmill for 1–2 weeks before experiments began.

The virtual-reality integration (VRI) task consisted of three components: a start cue, a cue-free running period, and an unmarked reward zone. Each trial began with a start cue presented on the corridor wall for 0.75 s. Animals could initiate running at any time after the previous trial ended, allowing the start cue and run onset to be temporally dissociated. The cue-free running period consisted of a 180-cm linear track with a constant gray corridor. Reward delivery (a drop of water via a spout) was triggered only if the animal actively licked within a 40-cm reward zone that began after the 180-cm track. Each trial ended with a 0.5-s inter-trial blackout. On rewarded trials, the blackout began immediately after reward delivery; on non-rewarded trials, it began when the animal exited the reward zone.

During initial training, water was delivered automatically when the animal reached the 180-cm position. After several days, the task was switched to the active-lick contingency, in which animals had to lick within the reward zone to receive water. Trial success was defined as trials in which a reward was triggered.

In dLight imaging experiments, the start cue was omitted from the VRI task to disambiguate run onset from cue onset as the trigger for time/distance estimation.

#### Pharmacology

Propranolol, prazosin, and SCH-23390 (all from Sigma-Aldrich, Massachusetts, USA) were dissolved in deionized water at a final concentration of 1 mM (0.296, 0.420, and 0.324 mg/mL, respectively).

Drug-infusion experiments via cannulae were performed using the following protocol. Animals were first recorded performing the task under normal conditions to establish pre-infusion behavioral performance. The task was then switched to passive mode (automatic water delivery at 180 cm) for drug delivery. Dummy cannulae were carefully removed, and the drug was infused (200 nL; 100 nL/min) through the cannulae targeting dorsal CA1 using a Hamilton syringe pump. Dummy cannulae were then replaced, and the task was switched back to active mode (lick-triggered reward delivery)^26^. Behavioral performance was recorded at 30 minutes post-infusion.

#### Optogenetics

Each optogenetic session consisted of a baseline block (40–50 trials), followed by a stimulation block (∼100 trials), and then a recovery block (40–50 trials).

To minimize the animal’s ability to distinguish stimulation and control trials, a constant “masking” light was delivered through a separate control optic fiber (470 nm LED; Thorlabs, New Jersey, USA) taped to the stimulation fibers, and remained on throughout the entire session.

For LC activation experiments without recording, and for LC activation combined with CA1 recording (electrophysiology or dLight imaging), bilateral stimulation was delivered at 12 Hz for 0.5 s (six pulses per train; 470 or 473 nm; 2–5 mW). The same blue light was used for hChR2 and ChrimsonR, avoiding red (∼590 nm) wavelengths to minimize PMT interference. Despite the lower sensitivity of ChrimsonR at ∼470 nm, stimulation reliably evoked CA1 dLight signals. Stimulation was triggered either at run onset (detected online when speed exceeded 10 cm/s for at least 0.2 s after the start cue), at mid-trial (120 cm), or at reward delivery.

For LC activation with LC recordings, and for LC–CA1 axon terminal activation with CA1 recordings, stimulation was delivered unilaterally through optic fiber(s) mounted on the probe (optrode, NeuroNexus), using the same stimulation protocol (0.5–2 mW).

For dLight imaging experiments with LC stimulation, light was delivered either at run onsets or manually via a control button in a customized MATLAB interface.

Stimulation and control trials were interleaved throughout the stimulation block, with one stimulation trial followed by two control trials (stimulation–control–control, repeating).

#### Electrophysiology

At least two days before acute electrophysiological recordings, mice were habituated to the head-fixed recording conditions. One day before recording, craniotomies were opened at pre-marked coordinates. Openings were temporarily protected with silicone gel (DOWSIL 3-4680) and covered with KwikSil or KwikCast (World Precision Instruments); these materials were removed immediately before probe insertion.

For LC recordings, a 32-channel single-shank silicon probe (A1x32-Poly3-5mm-25s-177, NeuroNexus) with an attached 50-μm-core optic fiber was used. The probe was lowered slowly into LC. In opto-tagging sessions, the actual recording depth was identified by delivering 5-ms tagging pulses at multiple depths below 2 mm. After reaching the target depth, the probe was allowed to stabilize for ≥5 min before recording. Optogenetic manipulations used the stimulation protocol described in the **Optogenetics** section. Opto- tagging was performed at the end of each session by delivering 60 light pulses (5 ms per pulse, with 5-s intervals in between) through the mounted optic fiber.

For CA1 recordings, a 64-channel, 6-shank silicon probe (buzsaki64sp, NeuroNexus; with or without optic fibers) was used. CA1 depth was guided by the expected position of the pyramidal layer, theta oscillation amplitude, the presence of sharp-wave ripples, and increased multi-unit activity and spike bursts^71^.

For both LC and CA1 recordings with optogenetics, stimulation followed the same protocols described in the **Optogenetics** section.

Signals were acquired with an Amplipex system, bandpass filtered (0.3 Hz–10 kHz), amplified (gain 400), and sampled continuously at 20 kHz.

#### Two-photon microscopy

After training, mice implanted with imaging cannulae were moved to the two-photon rig for habituation and subsequent imaging. Imaging was performed on a two-laser system consisting of a tunable femtosecond laser (InSight X3, Spectra-Physics) and a fixed-wavelength laser (Alcor 920, Spark Lasers). Fluorescence was acquired simultaneously in two channels (green and red).

For single-color green indicators (axon-GCaMP, dLight, GCaMP), imaging was performed with the fixed 920-nm laser. For dual-color imaging of dLight with jRGECO, the tunable laser was set to 1,000 nm. The excitation beam was delivered through a 16× water-immersion objective (N16XLWD-PF, 0.8 NA, 3.0 mm WD, Nikon). Emitted light was detected by a GaAsP photomultiplier tube (Hamamatsu Photonics) through bandpass filters for green (FF01-531/46; 508–554 nm) and red (FF01-625/90; 580–670 nm; Semrock). Image acquisition was controlled with ScanImage (MBF Bioscience) at 30 Hz (512 × 512 pixels; 3× optical zoom; 312 × 312 μm field of view). To align imaging to behavioral events, a copy of the frame trigger (“start-of-frame”) signal was routed to the behavioral control board.

Imaging was performed with laser power tuned to 20–40 mW, at depths ranging from 50– 200 μm below the glass surface.

For LC–CA1 axon imaging, axon-targeted GCaMP7b was expressed in LC neurons, and signals were recorded from LC axons in CA1 stratum radiatum and/or stratum pyramidale. For CA1 dLight imaging, dLight3.6 was expressed in CA1 neurons, and two-photon recordings were performed mainly from the CA1 stratum pyramidale. For experiments combining imaging with LC activation, tdTomato–ChrimsonR was expressed in LC, and optogenetic stimulation followed the protocol described in the Optogenetics section. To protect the PMT from stimulation light, the PMT shutters were closed during optical stimulation. For dopamine β-hydroxylase (DBH) inhibition, mice received nepicastat (25 mg/kg, i.p.; dose based on body weight; Selleck Chemicals). Experiments began 15 min after injection to allow systemic distribution and central uptake.

For CA1 dLight with jRGECO imaging, dLight3.6 and jRGECO1a were expressed in CA1 neurons, and two-photon recordings were performed mainly from the CA1 stratum pyramidale.

For CA1 GCaMP imaging with SCH-23390 local infusion, GCaMP8s was expressed in CA1 to monitor pyramidal neuron activity^52^. To allow local drug delivery, a small access hole was drilled through the imaging window using a 0.3-mm drill bit, and then the standard window implantation procedure was completed. Drug penetration was verified by infusing muscimol and observing the silencing of calcium activity. After each imaging recording, the imaging cannula was filled with sterile neurophysiological basal medium (BrainPhys; cat. #05791) and sealed with a plug to reduce inflammation and cell loss.

On each imaging day, a pre-infusion session with sterile saline was first recorded. Then saline was replaced with SCH-23390, prazosin, propranolol, or saline (vehicle control). A post-infusion session was recorded 20 min after infusion. Before the first drug day, animals were acclimated to the full infusion and imaging procedure with saline for at least 2 days.

### Quantification and statistical analysis

#### Behavioral analysis

Task-relevant behavioral signals (for example, running speed and licking) were extracted directly from the raw recordings. Derived measures (for example, first-lick time) were computed after segmenting the data into trials. Data were aligned to multiple task events, including start cue, run onset, and reward delivery. Run onset was defined as the start of the first sustained running bout after the previous trial had finished, when speed first exceeded 1 cm/s, provided that speed subsequently reached and remained above 10 cm/s for at least 0.3 s. First lick was defined as the first lick occurring after at least 1 s from run onset in the time domain, or 30 cm in the distance domain, to exclude licks carried over from the preceding trial.

#### Controls for locomotor kinematics

To test whether LC run-onset responses were explained by locomotor dynamics, trials were split within each session by initial speed (25–35 cm/s vs. 35–45 cm/s, 0–1 s window after run onset), initial acceleration (60–80 cm/s² vs. >80 cm/s², 0–1 s window after run onset), or mean speed (35–45 cm/s vs. 45–55 cm/s, over a trial). For each split, RON run-onset peak amplitudes were compared between the two groups.

#### Trial-start run onset versus spontaneous mid-trial run bouts

To assess whether LC run-onset activity was specific to the start of time/distance estimation rather than to the start of locomotion, spontaneous run onsets when animals paused in the middle of a trial and then resumed running were identified. Spontaneous run onsets were detected using the same criteria defined above for trial-start run onset. Run bouts occurring near the beginning of the trial (<50 cm) were excluded. Spontaneous run bouts were speed-matched to trial-start run bouts using the mean speed in the [-1, 1] s window relative to onset. RON firing rates were aligned to spontaneous run onsets and compared with trial-start run-onset responses.

#### Early versus late first-lick trials and speed matching

Early- and late-first-lick trials were defined for both LC and CA1 recordings. Trials were classified using fixed thresholds: early (<2.5 s) and late (2.5–3.5 s) relative to run onset. These thresholds captured the typical first-lick range; trials with first licks >3.5 s were often associated with failed trials or frequent or extended stopping.

To control for running speed, speed matching between early and late trials within each session was performed. Speed traces from 0–3.5 s after run onset were divided into seven 500-ms bins. For each bin, the mean and SD were computed separately for early and late trials. Trials were then filtered symmetrically: a late-first-lick trial was retained only if its binned speeds all fell within ±1 SD of the early-trial mean for each corresponding bin, and vice versa. The retained early and late trials after this reciprocal filtering formed the speed-matched sets. Sessions with fewer than 10 speed-matched trials in either condition were excluded from the CA1 analysis.

#### Pharmacology analyses

For pharmacological experiments, speed was compared using trial-averaged speed profiles. Early licking was quantified in the [30, 100] cm segment. Anticipatory licking was quantified in the [120, 180] cm segment, which immediately precedes the unmarked reward zone.

#### Optogenetic analyses

For LC optogenetic experiments, stimulation and interleaved control trials were analyzed separately and compared.

### Extracellular recording analysis

#### Preprocessing of LC recordings and opto-tagging

Spike sorting was performed offline using Kilosort^72^ and Klusters^73^. Units were manually curated, and those with clear refractory-period violations seen in the spike auto- correlogram were excluded.

For analysis, each neuron’s spike train was convolved with a Gaussian kernel (s.d. = 30 ms), aligned to behavioral events (start cue, run onset, and reward delivery), and averaged across trials to obtain an event-aligned mean firing-rate profile.

Opto-tagging was performed similarly to prior LC-tagging studies^31,32^. Units with mean firing rates > 20 Hz were excluded from tagging classification (mean firing rate > 10 Hz produces similar results). Among the remaining units, a neuron was classified as opto-tagged if it fired at least one spike within 10 ms after light-pulse onset on at least one-third of the 60 stimulation pulses.

#### Identification of putative LC Dbh^+^ and Dbh^−^ neurons

To identify additional catecholaminergic LC units beyond the opto-tagged population, Uniform Manifold Approximation and Projection (UMAP^30^) was used on spike auto- correlograms. Auto-correlograms (200-ms lag window, 1 ms bins) were smoothed (Gaussian kernel, s.d. = 0.5 ms), normalized, and z-scored, then embedded into two dimensions using UMAP. The 2D embedding separated units into two clusters, and all opto-tagged Dbh^+^ neurons fell within a single cluster. k-means clustering (k = 2) was applied to the UMAP embedding and putative Dbh^+^ neurons were defined as units assigned to the same cluster as the tagged Dbh^+^ neurons. To further restrict this set to physiological firing rates consistent with LC Dbh^+^ neurons, only units with a mean firing rate **<** 10 Hz were retained. Units assigned to the other cluster were classified as putative Dbh^−^ neurons.

Electrophysiological properties were compared across tagged Dbh^+^, putative Dbh^+^, and putative Dbh^−^ populations. Spike width was quantified as the full width at half maximum (FWHM) of the waveform after 10× temporal upsampling. Distributions of spike width and mean firing rate were compared between groups using Wilcoxon rank-sum tests.

#### Detection of LC run-onset neurons (RONs)

To identify neurons with significant run-onset firing, each unit’s run-onset-aligned mean firing-rate profile was compared to a null distribution generated by circularly shuffling spike timing within each trial (i.e., applying a random circular time shift to each trial’s spike train to preserve within-trial structure while disrupting alignment to run onset). For each cell, the trial-averaged firing-rate profile was computed after shuffling, and this procedure was repeated 5,000 times to obtain a distribution of shuffled mean profiles.

Within a ±0.5 s window around run onset, a significance threshold was defined as the 99th percentile of the shuffled mean profiles. A cell was classified as a run-onset neuron (RON) if its observed mean firing rate exceeded this threshold for at least 0.2 s continuously within the ±0.5 s run-onset window.

Throughout the paper, the amplitude of an RON’s run-onset phasic activity was quantified as the mean firing rate in the [-0.5, 0.5] s window relative to run onset.

For population heatmaps, LC neurons were ordered by their peak firing rate location. Heatmaps were normalized within each neuron (unless noted otherwise).

#### Generalized linear model for LC run-onset peak amplitude

To quantify which behavioral and neuronal variables explain trial-by-trial variability in the run-onset peak amplitude of LC run-onset–active neurons, we modeled peak amplitude on unstimulated trials (baseline trials plus optogenetic control trials).

For each neuron *n* and trial *t*, let *y_n_*_,*t*_ denote the run-onset peak amplitude (Hz). To stabilize variance and improve normality of residuals, we modeled the log-transformed response *z_n_*_,*t*_ = *log*(*y_n_*_,*t*_ + *ε*), *ε* = 10^−6^.

We constructed a prospective design matrix (predictors drawn from the current or previous trial), and z-scored predictors within session. The initial candidate set comprised 11 predictors.

There are four behavioral predictors: 1) Mean running speed on the previous trial, Speed*_t_*_−1_; 2) Lick count in the [-5, 0] s window relative to run onset, Licks*_t_*; 3) Reward-to-run interval on the current trial, *RRI_t_*; 4) Previous-trial timing error (“prediction error”), defined as the interval from first lick to reward delivery on trial *t* − 1, *Err_t_*_−1_.

There are seven neuronal predictors: 1) Baseline firing rate measured in five different pre-run windows (all ending at -2.5 s relative to run onset): [-30, -2.5] s, [-20, -2.5] s, [-15, - 2.5] s, [-10, -2.5] s, [-5, -2.5] s; 2) Previous-trial run-onset peak amplitude, *y_n_*_,*t*−1_; 3) Pre-onset firing rate in [-2.5, -1.5] s relative to run onset.

For each neuron, we fit a Gaussian linear regression of the log-transformed response:

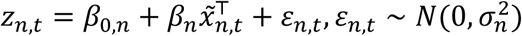

where ̃x*_n_*_,*t*_ is the z-scored predictor vector, and *ε_n_*_,*t*_ is Gaussian noise. Fits were obtained by maximum likelihood.

To assess the unique contribution of each predictor, we used a drop-one approach: for each predictor *j*, we refit a reduced model omitting *j* and computed the change in Akaike information criterion (AIC)

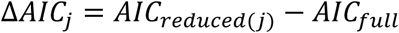

Positive Δ*AIC_j_* indicates worse fit when the predictor is removed.

To assess whether predictor contributions exceeded chance, we generated shuffled controls by randomly reassigning trial numbers (500 shuffles), refit the same models, and computed a null distribution of ΔAIC values. Based on these analyses, we retained three predictors that significantly improved fit: Speed*_t_*_−1_, RRI*_t_*, and baseline firing rate in [-5, -2.5] s. In addition, we included Err*_t_*_−1_ in our model, since recent literature has suggested that LC activity can reflect performance-related error signals^35^.

We therefore refit the final model using these four predictors. For each neuron and predictor *j*, we quantified incremental explained variance using drop-one-out *R*^2^:

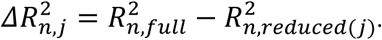

Significance was assessed against the permutation-based null models.

#### Correlation analysis for LC run-onset peak amplitude

We computed within-session correlations between trial-wise run-onset amplitude and reward-to-run interval. We split trials into “short” and “long” reward-to-run intervals (short: < 1.5 s; long: ≥ 1.5 s; criteria were selected based on the median of reward-to-run intervals across all behavioral sessions). We also computed session-wise correlations between (i) baseline firing rate (mean firing rate in [-1, -0.5] s relative to run onset) and reward-to-run interval, and (ii) baseline firing rate and run-onset firing rate.

#### Preprocessing of CA1 recordings

CA1 spike sorting was performed as for LC recordings. Putative pyramidal neurons were identified based on spike waveform features and mean firing rate (0.15–7 Hz). Spike trains were convolved with a Gaussian kernel (s.d. = 30 ms), aligned to behavioral events, and trial-averaged firing-rate profiles were computed.

#### Identification of PyrUp and PyrDown neurons

PyrUp and PyrDown pyramidal neurons^26^ were classified using a firing-rate ratio (R) computed from run-onset–aligned activity:

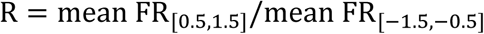

Neurons with R > 1.5 were classified as PyrUp, and neurons with R < 0.67 were classified as PyrDown.

For population heatmaps, pyramidal neurons were ordered by FR ratio. Heatmaps were normalized within each neuron (unless noted otherwise).

#### Two-photon imaging analysis Preprocessing of two-photon movies

Raw movies were first corrected for x–y motion using the Suite2p registration pipeline. Motion estimates were computed from a single reference channel and applied to both channels to preserve channel alignment^74^. The reference channel was selected based on the experimental configuration: (1) LC–CA1 axon-GCaMP imaging used the green channel; (2) CA1 dLight imaging during optogenetic LC stimulation, as well as CA1 dLight imaging in Dbh-Cre::Ai14 mice, used the tdTomato channel (red); (3) dual-color dLight + jRGECO imaging used the green channel as the reference; and (4) CA1 GCaMP imaging during D_1_- like receptor blockade, performed in Drd1-Cre::Ai14 mice that express tdTomato in Drd1^+^ neurons, used the red channel as the reference.

### ROI detection for two-photon imaging

#### LC–CA1 axon-GCaMP imaging

We identified candidate axonal ROIs using Suite2p functional ROI detection applied to the GCaMP channel. Then we manually curated ROIs to (i) merge fragments likely belonging to the same axon and (ii) remove spurious detections.

#### CA1 dLight imaging during optogenetic LC stimulation

We identified LC axons in the field of view to quantify how dLight responses vary with distance to axons. Because LC axons were labeled with tdTomato (a static anatomical marker with no activity-dependent fluctuations), we did not use the Suite2p activity-based ROI detection. Instead, we developed a custom graphical user interface (GUI) to segment LC axons from the tdTomato reference images (red channel).

To facilitate axon detection, we first preprocessed each reference image to enhance thin, fiber-like structures and reduce uneven background. Specifically, we applied a white top- hat filter to emphasize narrow bright features, followed by adaptive histogram equalization (CLAHE; OpenCV) to improve local contrast. We then extracted candidate axon ROIs using the Maximally Stable Extremal Regions (MSER) algorithm^75^. Next, we filtered candidate regions using geometric criteria (for example, minimum area, aspect ratio, eccentricity, solidity, and circularity index) to select fiber-like structures and exclude round or punctate objects.

We then displayed detected candidate ROIs on the reference image in the custom GUI, and manually curated ROIs by accepting, deleting, or merging candidates to produce the final set of LC-axon ROIs used in downstream analyses.

#### CA1 dLight imaging in Dbh-Cre::Ai14 animals

For dLight imaging during behavior, we included only recordings with stable task performance and minimal movement-related artifacts. We assessed task performance using lick selectivity, maximum running speed, and maximum trial duration. To quantify movement artifacts, we computed the correlation between the whole-field dLight signal and the animal’s running speed. We retained recordings in which the dLight-speed correlation was below 0.3 (indicating low motion contamination) for analysis.

We detected and curated LC axons using the same procedure as in the dLight imaging with optogenetic LC stimulation experiments. We partitioned the 512×512 pixel field of view into 32×32 bins (∼10×10 μm per bin), and termed bins containing tdTomato-positive LC axon pixels “axon bins”. Within each axon bin, the tdTomato-positive pixels that also showed dLight expression defined the axon-containing region (ACR). We then extracted dLight signal from each ACR. To adaptively detect dLight expression pixels with different morphology and intensity, we first smoothed the reference dLight field of view (FOV) with a 2D Gaussian filter (s.d. = 1.5 pix). Then we computed a local “uniformity” metric in non-overlapping 10×10 pixel blocks:

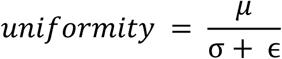

where *μ* is the local region mean, σ is the standard deviation of the local region, ɛ = 1e-6.

Using this metric, we separated the FOV into two compartments: (i) a **cell layer**, containing somata with membrane-localized dLight expression, and (ii) a **non-cell layer**, containing dendrites/axons. For each ACR, we defined the neuropil within a square region centered on the ACR: we excluded a 3-pixel buffer around the ACR, and then expanded the square outward in 5-pixel increments until the neuropil mask contained 350 pixels. In addition, we defined the local dLight background region as dLight-positive pixels within a 3-pixel binary dilation ring surrounding the ACR (excluding the ACR itself) in the same bin.

#### CA1 dLight and jRGECO dual-color imaging

We detected ROIs using the pre-trained cyto model in Cellpose^76^. The neuropil and dLight background regions were defined as described above for **CA1 dLight imaging in Dbh-Cre::Ai14 animals**. To restrict analyses to somatic ROIs and exclude elongated axonal or dendritic masks, we applied anatomical filters requiring each ROI to contain 100–600 pixels and to have an aspect ratio < 3.

#### CA1 GCaMP imaging with D_1_-like receptor blockade

We detected ROIs using Suite2p, and selected somatic ROIs using the criteria described above for **CA1 dLight and jRGECO dual-color imaging**.

### Movement artifact correction using linear regression

#### CA1 dLight imaging in Dbh-Cre::Ai14 animals

For each axon bin, we extracted dLight and tdTomato fluorescence from its ACR. We also extracted dLight neuropil and local background signals. dLight signals were first neuropil-corrected.

To further reduce potential motion-related artifacts, we fit a linear regression model using the dLight background signal and the axonal tdTomato signal as regressors. The model was fit separately for each trial segment spanning [-90, 120] frames around run onset:

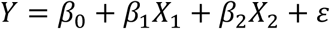

where *Y* is the neuropil-corrected dLight signal, *X*_1_ is the dLight background signal, and *X*_2_ is the tdTomato signal.

#### CA1 dLight and jRGECO dual-color imaging

We extracted jRGECO ROI and neuropil signals in Suite2p using the masks described above. For each jRGECO ROI, we extracted dLight fluorescence from the subset of ROI pixels that expressed dLight. Signals were neuropil-corrected.

We classified frames in which the jRGECO signal fell below median + 2 × robust SD as baseline (non-active) periods. Residual fluctuations during these periods were attributed to motion artifacts and used as a regressor. We then used linear regression to remove residual motion artifacts from the neuropil-corrected dLight signal, with two regressors: (i) the dLight background signal and (ii) the jRGECO baseline signal. We fit the regression separately for each trial segment ([-90, 120] frames relative to run onset).

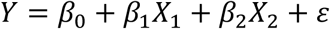

where *Y* is the neuropil-corrected dLight signal, *X*_1_ is the dLight background signal, and *X*_2_ is the jRGECO baseline signal.

For dLight signals in both Dbh-Cre::Ai14 and dual-color experiments, we computed the predicted artifact signal from the regression and subtracted it from the neuropil-corrected dLight trace:

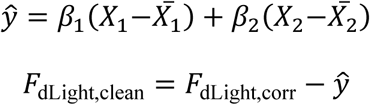

where *̂y* is the estimated artifact contribution from *X*_1_ and *X*_2_.

#### Fluorescence normalization (ΔF/F)

Fluorescence was converted to ΔF/F in two steps. First, a rolling baseline *F*_0_ was defined as the n-th percentile of raw fluorescence values (F) within a 300 s sliding window. Second, ΔF/F was calculated using

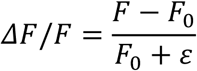

where ε = 10^−9^ or 10^−8^, added to prevent division by zero in dark regions. n = 20 for GCaMP/jRGECO signals, and 15 for dLight signals.

#### LC–CA1 axon-GCaMP signal analysis

The ΔF/F trace was aligned to different behavioral events. For trial-wise analyses, ΔF/F traces of each ROI were smoothed with a Gaussian kernel (s.d. = 3 frames). Run-onset- active axons were detected using the same peak-detection procedure as for LC RONs.

#### Comparison of calcium responses between DA-Up and non-DA-Up neurons

To compare the run-onset calcium response between DA-Up and non-DA-Up neurons, the ΔF/F was baseline-corrected by subtracting the mean of ΔF/F within the [-0.5, 0] s window before run onset.

### Analysis for CA1 dLight imaging with LC optogenetic stimulation

#### Pixel-wise dLight signal analysis

The motion-registered dLight movie was spatially smoothed with a Gaussian kernel (s.d. = 1 pixel). The movie was then aligned to the time of each optogenetic stimulation train.

For each stimulation epoch, a pixel-wise response index (RI) was computed to summarize the stimulation-evoked change in dLight signal:

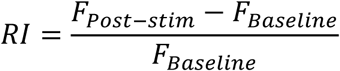

where *F*_Baseline_ was the mean raw fluorescence in the pre-stimulation baseline window (- 1.00 to 0 s relative to stimulation onset), and *F*_Post−stim_ was the mean raw fluorescence in the post-stimulation window (for example, 0 to 1.00 s relative to stimulation offset). RI was computed independently for every pixel and every stimulation epoch. To prevent numerical instability in pixels with very low baseline fluorescence, RI values were capped at 10.

To visualize the post-stimulation time course of the spatial pattern, the same computation was repeated using *F*_Post−stim_ measured in 0.1 s non-overlapping bins spanning 0–4 s after stimulation offset (40 bins total), while keeping the *F*_Baseline_ as previously defined. This produced a 3D RI array (512 × 512 × 40) per stimulation epoch describing the evolution of pixel-wise dLight responses over time. The median of this 3D RI array was then computed across stimulation epochs within each recording for subsequent analyses.

#### LC-axon-pixel-based analysis of dLight signal

LC-axon ROIs were first delineated in the tdTomato reference channel using a custom graphical interface (see the **ROI detection for two-photon imaging** above). Axon ROIs were classified as responding or non-responding based on their pixel-by-pixel stimulation-evoked RI, computed for each stimulation epoch and then averaged across epochs. A responding ROI was required to: 1) have a mean RI across pixels greater than zero (one-sample t-test, p < 0.01), and 2) have a minimum RI across pixels of at least 0.10.

For each session, all responding axon ROIs were combined to generate a single binary mask of LC-axon pixels (responding axon pixels = 1, otherwise = 0). We then computed the RI time course for LC axons by averaging the stimulation-epoch-averaged 3D RI array across all axon pixels in the mask at each time bin.

In parallel, a background mask was defined as all pixels at least 9 pixels (∼5.48 μm) from the nearest LC axon ROI, and the corresponding background RI time course was computed in the same way.

#### Temporal decay of dLight signal

Decay kinetics of the RI time course were estimated by fitting an exponential function to each RI time course:

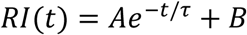

where *A* is the response amplitude, *B* is the offset, and *τ* is the decay time constant.

#### Spatial decay of dLight signal

To quantify how RI decreases with distance from LC axons, we measured RI in concentric 1-pixel-wide “rings” surrounding LC-axon ROIs. We iteratively dilated each axon ROI in 1-pixel steps (up to 20 pixels). At each step, we defined a ring mask as the set of pixels newly added by that dilation, yielding a 1-pixel-wide ring at a defined distance from the axon ROI.

For each session, we combined rings at the same distance across responding axon ROIs to form a single binary mask per distance. We then computed the mean RI of ring pixels in the [0, 1] s window after stimulation offset to generate an RI-versus-distance profile. Finally, we fit an exponential decay function to this spatial profile to obtain a session-level spatial decay constant, which was used for group-level analyses.

#### Control for dLight expression gradients

To test whether the heterogeneous RI across the field can be explained by spatial differences in dLight expression, we regressed baseline dLight fluorescence (mean dLight intensity across the session for each pixel) against the corresponding RI values. This analysis was performed both across the full field of view and restricted to LC-axon pixels.

### Analysis for CA1 dLight imaging in Dbh-Cre::Ai14 animals

#### Identification of run-onset DA-Up-ACRs

To identify ACRs with significant dopamine increases at run onset (DA-Up-ACRs), we computed two metrics from the regression-corrected dLight signal of each ACR.

We first defined a response amplitude as the difference in mean Δ*F*/*F* between a post–run-onset window and a pre–run-onset baseline window:

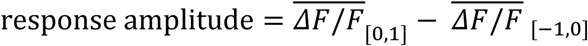

Second, we computed Cohen’s *d* to quantify the effect size, defined as response amplitude normalized by the pooled standard deviation.

An ACR was classified as a DA-Up-ACR if it met two criteria: (i) Cohen’s *d* > 0.05 and (ii) its response amplitude exceeded the 95th percentile of a shuffled null distribution.

#### Temporal and spatial decay of dLight signal

To quantify the temporal decay of the dLight signal, we fit the trial-averaged trace of each DA-Up-ACR with an exponential decay function, using the same procedure as in **Analysis for CA1 dLight imaging with LC optogenetic stimulation**. To estimate spatial decay, we computed response amplitudes from progressively dilated ACR masks using the same response-amplitude definition applied to ACRs, and fit an exponential to the resulting amplitude-versus-distance profile.

### Analysis for CA1 dLight and jRGECO dual-color imaging

#### PyrUp and PyrDown classification for jRGECO ROIs

To facilitate comparison with electrophysiological recordings, we limited our analyses to somatic ROIs (see ROI mask criteria) that exhibited detectable calcium activity during the session. A fluorescence-trace skewness > 0.8 was selected based on visual inspection of individual fluorescence traces to distinguish active neurons from inactive neurons.

We then classified active neurons as PyrUp or PyrDown using the ratio of mean jRGECO Δ*F*/*F* after versus before run onset:

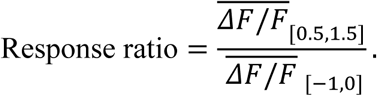

ROIs with response ratio > 1.12 were classified as PyrUp, whereas ROIs with response ratio < 1/1.12 were classified as PyrDown. We selected the threshold to approximately match the percentages of PyrUp/PyrDown neurons identified in electrophysiological experiments.

#### CA1 GCaMP imaging with D_1_-like receptor blockade

The same classification procedure as described above was applied to CA1 GCaMP recordings during D_1_-like receptor blockade, using identical skewness and response-ratio thresholds.

### Computational modeling

#### Modeling method

To test whether PyrUp-specific modulation could arise from activity-dependent DA sensitivity rather than cell-type-specific targeting, we built an LC–dopamine–CA1 rate model.

##### Architecture

The model contained 1,000 CA1 neurons, each receiving three time-varying inputs: (i) a run-related input that activated rapidly at run onset and decayed gradually toward reward, R(t), (ii) a reward-related input that was suppressed at run onset and ramped up toward reward, W(t), and (iii) a dopamine input derived from a phasic LC signal at run onset, D(t). Each neuron was assigned a random baseline excitability, run-related input weight, and reward-related input weight, drawn independently from Gaussian distributions.

##### LC-derived dopamine signal

The phasic LC signal was modeled as a Gaussian transient (mean = 0 s, s.d. = 0.2 s), convolved with a causal exponential kernel (τ = 1.6 s, roughly consistent with the temporal decay measured in dLight experiments) to generate an LC-derived dopamine release waveform, *D_release_*(*t*). This waveform was then delayed by δ*_DA_* = 0.35 s to approximate the latency between DA release and D_1_-like receptor activation in CA1:

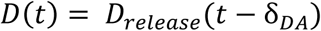

##### Activity-dependent dopamine modulation

Thirty percent of neurons were randomly designated as dopamine-targeted, approximating the proportion observed in dLight imaging experiments. Each dopamine-targeted neuron was assigned a random dopamine sensitivity parameter drawn from a uniform distribution on [0, 1]. The delayed dopamine signal, *D*(*t*), was further scaled by an activity-dependent gain that depended on the neuron’s firing rate at the previous time step, *r_i_*(*t* − *dt*) (**Fig. 5A**, inset), such that neurons with higher ongoing activity received stronger dopamine amplification.

##### Firing rate computation

For each neuron *i*, the non-DA input was:

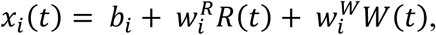

where *b_i_* was the neuron’s baseline excitability, 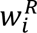 and 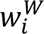 were cell-specific run-related and reward-related input weights.

The full input including dopamine was:

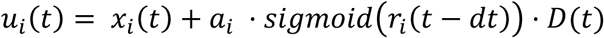

where *a_i_* is the neuron-specific dopamine sensitivity (0 for non-targeted neurons, drawn from [0, 1] for targeted neurons), *r_i_*(*t*) is the neuron’s instantaneous firing rate, and *D*(*t*) is the delayed dopamine signal. This input was passed through a first-order low-pass filter (leaky integrator) with a neuron-specific time constant *τ_i_*, and the resulting latent state was transformed through a softplus nonlinearity (*β* = 2) with a maximum rate of 20 Hz to produce the final firing rate.

##### Simulation

The model was simulated from −1 to 6 s relative to run onset with a 10-ms time step. Neurons were classified as PyrUp (R > 1.5) or PyrDown (R < 0.67) using the same firing-rate ratio as in the experimental analyses.

##### In silico experiments

We performed three simulations corresponding to the main experimental manipulations. First, we compared DA-targeted and non-DA-targeted neurons under baseline conditions (corresponding to **Fig. 5B**). Second, we increased the amplitude of the phasic LC transient to simulate optogenetic LC activation **(Fig. 5C**). Third, we removed the dopamine drive while leaving run-related and reward-related inputs unchanged to simulate D_1_-like receptor blockade (f**ig. S11**). Each simulation was run 50 times with independent random seeds, each generating a new population of 1,000 neurons.

#### Statistics and reproducibility

Data are reported as mean ± SEM unless stated otherwise; when medians are reported, they are accompanied by the interquartile range (IQR). Statistical tests were two-sided. Nonparametric Wilcoxon signed-rank (paired) or rank-sum (unpaired) tests were used by default. Paired or unpaired t-tests were used in cases where the data distribution was approximately symmetric and did not show extreme outliers. When multiple comparisons were performed within a single analysis (for example, across distance bins in the dLight spatial-dilation analysis), p-values were adjusted using Bonferroni correction.

## Code availability

Most custom scripts for analysis are available on GitHub (https://github.com/dinghaoluo/lc-ca1-project; https://github.com/yuxi-yuxi/code_mpfi_Jingyu). The remaining scripts are available upon reasonable request.

## Data availability

Data corresponding to all main figures are to be deposited at an open-source repository upon publication.

## Acknowledgments

We thank M. Klement, N. Daniel, and the MPFI machine shop for fabricating mechanical parts for the experimental setups; J. Wells, J. Poole-Capron, and the MPFI Animal Resources Center (ARC) for animal care; J. Yu and the MPFI molecular core for virus production and verification; and Y. Long and the MPFI microscopy core for support with imaging equipment. This work was funded by the Max Planck Society and the Max Planck Foundation.

## Author contributions

Y.W. and D.L. conceived the project. D.L., J.C., and Y.W. designed the experiments. D.L. and J.C. performed the experiments. D.L., J.C., and Y.W. performed data analysis. R.H. provided data for CA1 electrophysiology recordings. L.T. provided optimized dLight sensors; D.L., J.C., and Y.W. wrote the manuscript with input from all authors. Y.W. acquired funding, provided resources, supervised personnel, and led the project.

