## Supplemental material for "Spatially confined dopamine from locus coeruleus axons selectively shapes hippocampal dynamics and action timing"

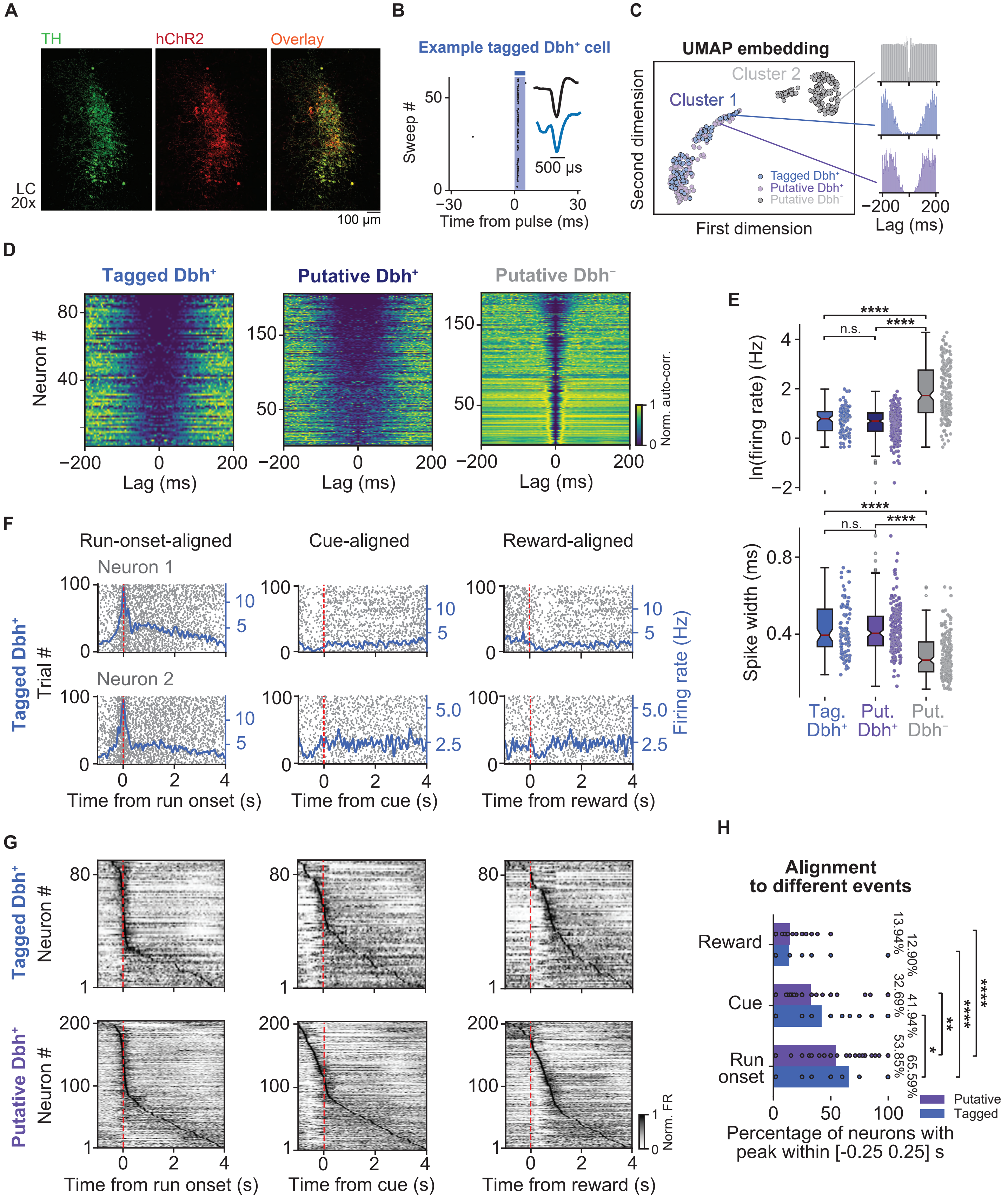
A
TH
hChR2
Overlay
LC
20x
100 µm
B
Example tagged Dbh⁺ cell
Sweep #
500 µs
Time from pulse (ms)
C
UMAP embedding
Cluster 2
Cluster 1
Second dimension
First dimension
Tagged Dbh⁺
Putative Dbh⁺
Putative Dbh⁻
Lag (ms)
D
Tagged Dbh⁺
Putative Dbh⁺
Putative Dbh⁻
Neuron #
Lag (ms)
Norm. auto-corr.
E
ln(firing rate) (Hz)
Spike width (ms)
n.s.
****
Tag. Dbh⁺
Put. Dbh⁺
Put. Dbh⁻
F
Run-onset-aligned
Cue-aligned
Reward-aligned
Neuron 1
Neuron 2
Tagged Dbh⁺
Trial #
Firing rate (Hz)
Time from run onset (s)
Time from cue (s)
Time from reward (s)
G
Tagged Dbh⁺
Putative Dbh⁺
Neuron #
Norm. FR
Time from run onset (s)
Time from cue (s)
Time from reward (s)
H
Alignment to different events
Reward
Cue
Run onset
13.94%
12.90%
32.69%
41.94%
53.85%
65.59%
Putative
Tagged
Percentage of neurons with peak within [-0.25 0.25] s

**Fig. S1 Opto-tagging, classification, and event alignment of LC Dbh+ neurons, related to Fig. 1**

(**A**) Representative histology from LC showing $Dbh^+$ neurons immunostained for tyrosine hydroxylase (TH; green) and virally expressed hChR2 (red); right, overlay. (**B**) Example optogenetically tagged $Dbh^+$ neuron. Raster plot shows light-evoked spikes across repeated optical pulses. Inset shows the average spike waveforms, comparing light-evoked waveform (blue) with spontaneous waveform (black). (**C**) Classification of recorded LC units. Left: UMAP embedding of spike auto-correlograms showing tagged $Dbh^+$ (blue), putative $Dbh^+$ (purple), and putative $Dbh^-$ neurons (gray). Tagged $Dbh^+$ and putative $Dbh^+$ neurons co-localized within Cluster 1. Representative auto-correlograms from example neurons are shown on the right. (**D**) Heatmaps of normalized spike auto-correlograms of tagged $Dbh^+$ (left), putative $Dbh^+$ (middle), and putative $Dbh^-$ neurons (right). (**E**) Summary of firing rates (top; ln-transformed) and spike widths (bottom) for tagged $Dbh^+$, putative $Dbh^+$, and putative $Dbh^-$ neurons. Each dot represents one neuron (natural log of firing rate: tagged $Dbh^+$ vs. putative $Dbh^+$, $p = 0.27$; tagged $Dbh^+$ vs. putative $Dbh^-$, $p = 5.5e-20$; putative $Dbh^+$ vs. putative $Dbh^-$, $p = 8.08e-34$; spike width: tagged $Dbh^+$ vs. putative $Dbh^+$, $p = 0.98$; tagged $Dbh^+$ vs. putative $Dbh^-$, $p = 3.46e-16$; putative $Dbh^+$ vs. putative $Dbh^-$, $p = 2.54e-26$; all Wilcoxon rank-sum tests). (**F**) Top and bottom rows show two example tagged $Dbh^+$ neurons, each aligned to trial-start run onset (left), start cue (middle), and reward delivery (right). Gray, single-trial rasters; blue, trial-averaged firing rate. (**G**) Population heatmaps of trial-averaged firing rates for tagged (top) and putative (bottom) $Dbh^+$ neurons aligned to run onset (left), start cue (middle), or reward delivery (right). Neurons are ordered by peak time. (**H**) Quantification of event alignment. Bars indicate the percentage of neurons with significant peak firing within ±0.25 s of the indicated event for tagged and putative $Dbh^+$ populations. Each dot represents one recording (tagged $Dbh^+$: run vs. cue, $p = 1.43e-02$; run vs. reward, $p = 9.69e-06$; putative $Dbh^+$: run vs. cue, $p = 1.56e-03$; run vs. reward, $p = 2.46e-06$; 8 animals, 39 recordings for tagged $Dbh^+$ and 35 recordings for putative $Dbh^+$; all Wilcoxon signed-rank tests).

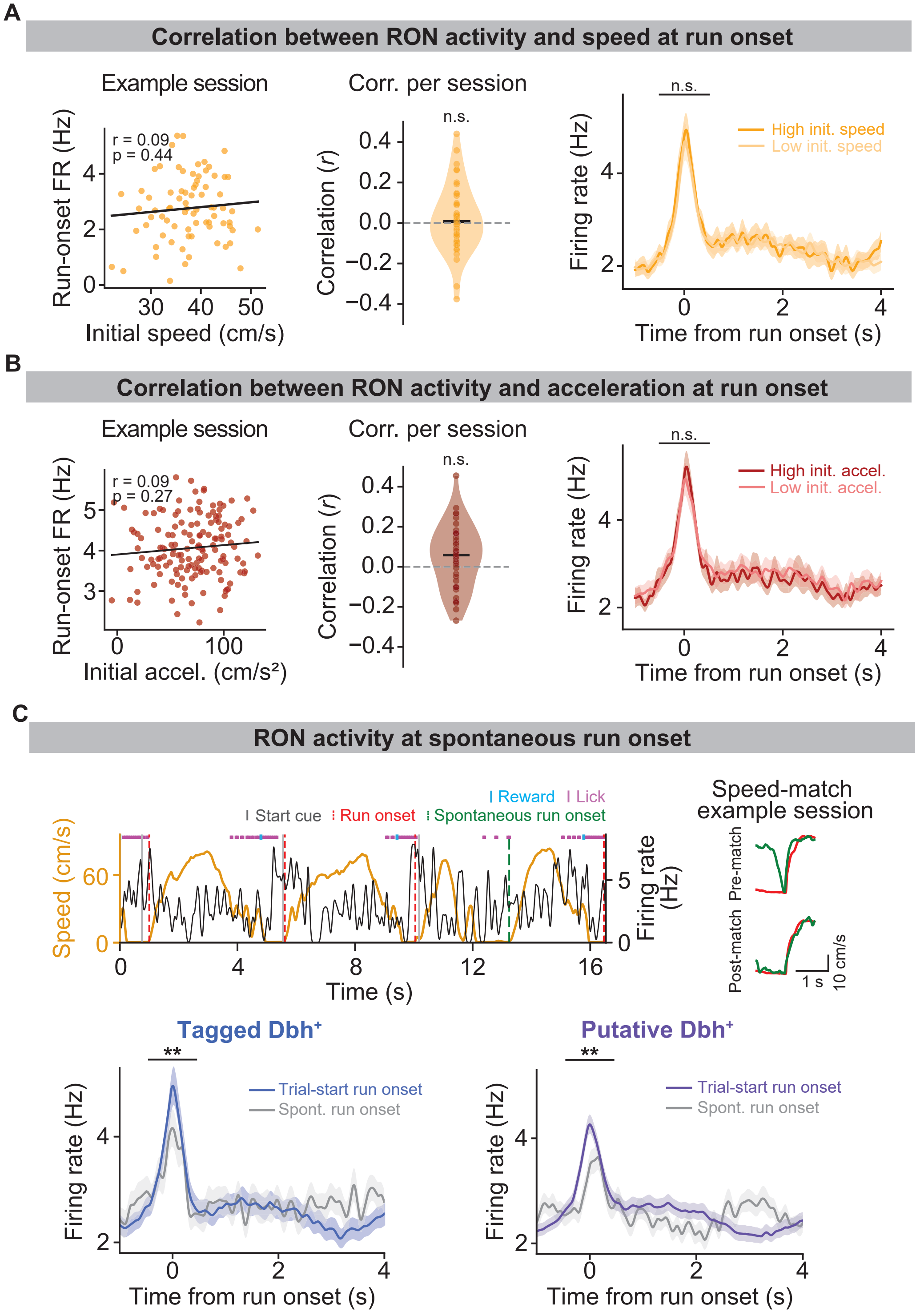
A
Correlation between RON activity and speed at run onset
Example session
r = 0.09
p = 0.44
Run-onset FR (Hz)
Initial speed (cm/s)
Corr. per session
n.s.
Correlation (r)
n.s.
High init. speed
Low init. speed
Firing rate (Hz)
Time from run onset (s)
B
Correlation between RON activity and acceleration at run onset
Example session
r = 0.09
p = 0.27
Run-onset FR (Hz)
Initial accel. (cm/s²)
Corr. per session
n.s.
Correlation (r)
n.s.
High init. accel.
Low init. accel.
Firing rate (Hz)
Time from run onset (s)
C
RON activity at spontaneous run onset
I Reward
I Lick
Start cue
Run onset
Spontaneous run onset
Speed (cm/s)
Firing rate (Hz)
Time (s)
Speed-match example session
Pre-match
Post-match
1 s
10 cm/s
Tagged Dbh+
**
Trial-start run onset
Spont. run onset
Firing rate (Hz)
Time from run onset (s)
Putative Dbh+
**
Trial-start run onset
Spont. run onset
Firing rate (Hz)
Time from run onset (s)

**Fig. S2 Run-onset LC phasic responses are not explained by locomotor kinematics, related to Fig. 1**

(**A**) RON firing rate versus running speed at run onset. Left, example session showing the relationship between run-onset speed (mean speed [0, 1] s) and run-onset RON firing rate. Each orange dot is one trial. Black line: linear fit. Middle, distribution of Pearson correlation coefficients computed per session (median = 0.01, IQR = [−0.09, 0.11]; p = 0.60; one-sample *t*-test against 0). Right, mean RON firing-rate profiles split by low versus high run-onset speed trials (low speed, median = 3.13 Hz, IQR = [2.66, 3.89] Hz; high speed, median = 3.15 Hz, IQR = [2.60, 4.08] Hz; p = 0.75; 7 animals, 20 recordings; Wilcoxon signed-rank test). (**B**) Same as (**A**), but for the relationship between run-onset firing rate and run-onset acceleration (correlation with run-onset acceleration: median = 0.06, IQR = [−0.08, 0.17]; p = 0.09; one-sample *t*-test against 0; run-onset firing rate: low acceleration, median = 3.33 Hz, IQR = [2.63, 4.26] Hz; high acceleration, median = 3.19 Hz, IQR = [2.81, 4.26] Hz; p = 0.90; Wilcoxon signed-rank tests). (**C**) RON firing rate at trial-start run onset and spontaneous run onset. Top left: example consecutive trials showing running speed (orange) and firing rate of an RON (black). Gray: start cue onset; red: trial-start run onset; green: spontaneous run onsets. Top right: example recording illustrating trial-averaged running speed before and after speed-matching procedure for trial-start versus spontaneous run onsets. Bottom: mean firing rate profiles aligned to run onset for tagged (left) and putative $Dbh^+$ neurons (right), comparing trial-start run onsets (colored) to spontaneous run onsets (gray) (tagged $Dbh^+$: trial-start run onsets, median = 3.15 Hz, IQR = [2.69, 4.10] Hz; spontaneous run onsets, median = 2.97 Hz, IQR = [2.14, 3.87] Hz; p = 6.09e-03; putative $Dbh^+$: trial-start run onsets, median = 3.15 Hz, IQR = [2.43, 4.24] Hz; spontaneous run onsets, median = 2.74 Hz, IQR = [1.85, 3.96] Hz; p = 5.46e-03; 7 animals, 20 recordings; both Wilcoxon signed-rank tests).

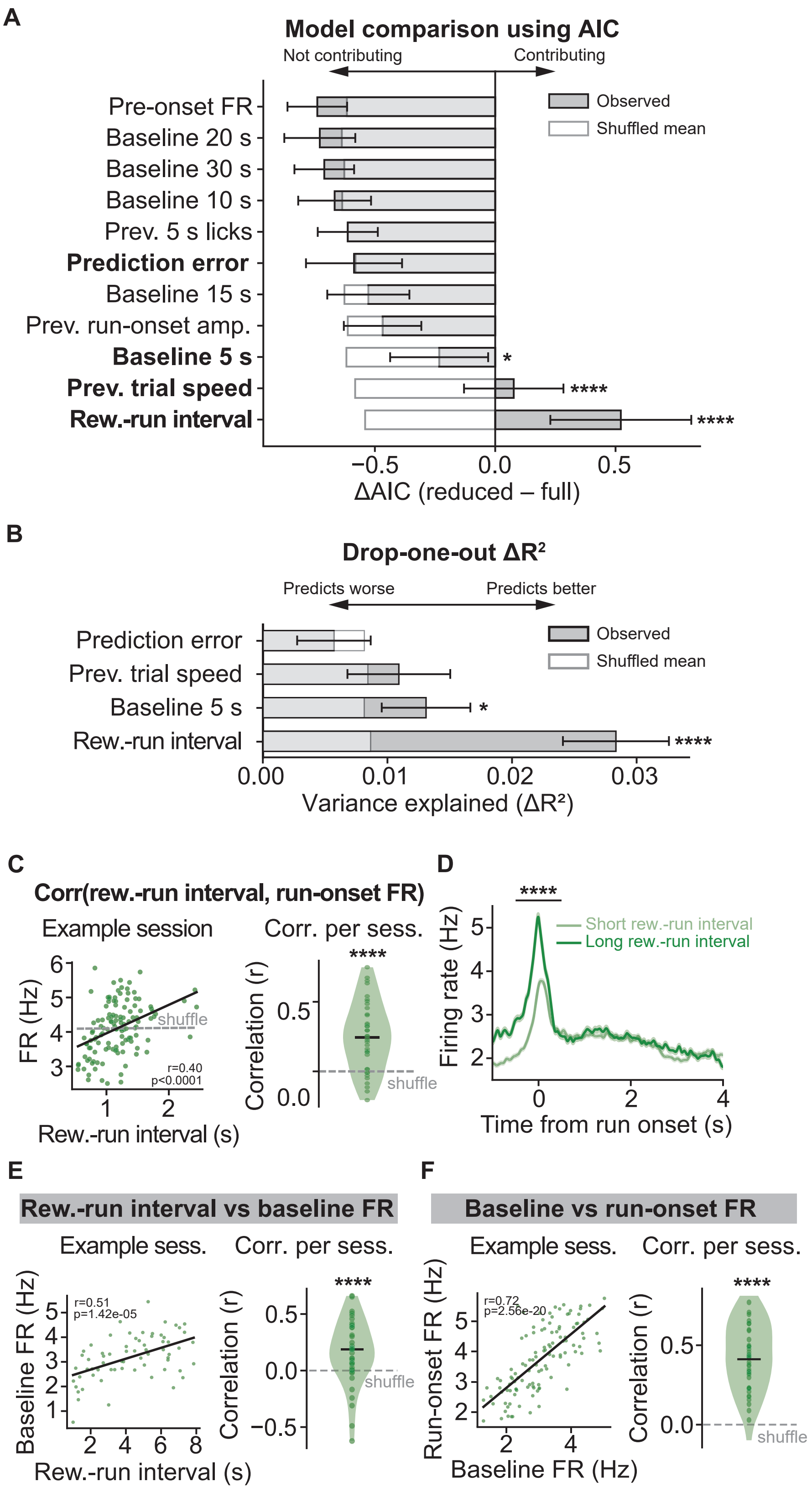
A
Model comparison using AIC
Not contributing
Contributing
Observed
Shuffled mean
Pre-onset FR
Baseline 20 s
Baseline 30 s
Baseline 10 s
Prev. 5 s licks
Prediction error
Baseline 15 s
Prev. run-onset amp.
Baseline 5 s
Prev. trial speed
Rew.-run interval
*
****
****
−0.5
0.0
0.5
ΔAIC (reduced – full)
B
Drop-one-out ΔR²
Predicts worse
Predicts better
Observed
Shuffled mean
Prediction error
Prev. trial speed
Baseline 5 s
Rew.-run interval
*
****
0.00
0.01
0.02
0.03
Variance explained (ΔR²)
C
Corr(rew.-run interval, run-onset FR)
Example session
Corr. per sess.
FR (Hz)
shuffle
r=0.40
p<0.0001
Rew.-run interval (s)
Correlation (r)
****
shuffle
D
****
Short rew.-run interval
Long rew.-run interval
Firing rate (Hz)
Time from run onset (s)
E
Rew.-run interval vs baseline FR
Example sess.
Corr. per sess.
r=0.51
p=1.42e-05
Baseline FR (Hz)
Rew.-run interval (s)
Correlation (r)
****
shuffle
F
Baseline vs run-onset FR
Example sess.
Corr. per sess.
r=0.72
p=2.56e-20
Run-onset FR (Hz)
Baseline FR (Hz)
Correlation (r)
****
shuffle

**Fig. S3 Reward-to-run interval predicts RON firing rate at run onset, related to Fig. 1**

(**A**) Model comparison for candidate predictors of RON run-onset firing rate. For each predictor (listed on the y-axis), we compared a full GLM containing all predictors to a reduced model with that predictor removed. Bars show $\Delta AIC = AIC_{reduced} - AIC_{full}$ for the observed data (dark) and a shuffled control (light). AIC: Akaike information criterion. Predictors carried forward to the simplified model in (**B**) are shown in bold. (**B**) Unique variance explained by each retained predictor in the simplified model, quantified as the drop-one-out $\Delta R^2$. (**C**) Left, example recording showing the relationship between reward-to-run intervals and RON firing rate at run onset. Each green dot is one trial. Black line: linear fit; gray dashed line: shuffle. r and p are from a Pearson correlation for that recording. Right, distribution of Pearson correlation coefficients computed per recording, compared with a shuffled distribution (median r = 0.21, IQR = [0.04, 0.37]; p = 7.88e-07; 7 animals, 20 recordings; Wilcoxon signed-rank test). (**D**) Mean firing rate profiles of RONs separated by short (light green) vs. long (dark green) reward-to-run intervals (firing rate at run onset: short reward-run intervals, median = 2.77 Hz, IQR = [1.90, 3.94] Hz; long reward-run intervals, median = 3.25 Hz, IQR = [2.31, 4.31] Hz; p = 2.52e-09; Wilcoxon signed-rank test). (**E**) Correlation between reward-to-run interval and RON baseline firing rate ([-1, -0.5] s). Left, example recording; right, distribution of Pearson correlation coefficients computed per recording. Each dot represents one recording (median = 0.19, IQR = [0.01, 0.38]; p = 4.13e-04; 7 animals, 20 recordings; Wilcoxon signed-rank test). (**F**) Correlation between RON baseline firing rate and RON firing rate at run onset. Left, example recording; right, distribution of Pearson correlation coefficients computed per recording (median = 0.41, IQR = [0.27, 0.60]; p = 1.09e-12; Wilcoxon signed-rank test).

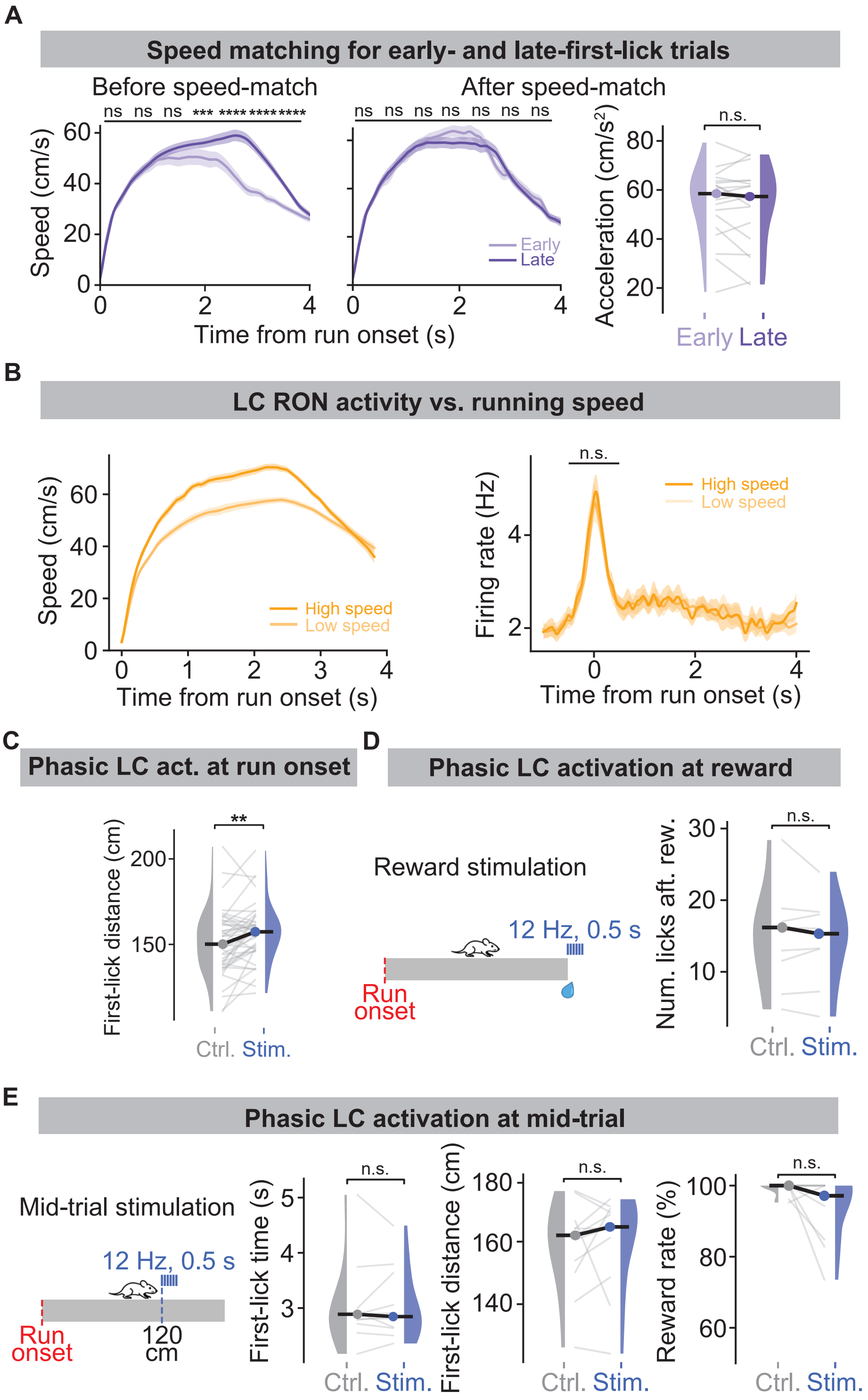
A
Speed matching for early- and late-first-lick trials
Before speed-match
After speed-match
ns ns ns *** ************
ns ns ns ns ns ns ns
Speed (cm/s)
Time from run onset (s)
Early
Late
Acceleration (cm/s²)
n.s.
Early Late
B
LC RON activity vs. running speed
Speed (cm/s)
High speed
Low speed
Time from run onset (s)
n.s.
Firing rate (Hz)
High speed
Low speed
Time from run onset (s)
C
Phasic LC act. at run onset
**
First-lick distance (cm)
Ctrl. Stim.
D
Phasic LC activation at reward
Reward stimulation
12 Hz, 0.5 s
Run onset
n.s.
Num. licks aft. rew.
Ctrl. Stim.
E
Phasic LC activation at mid-trial
Mid-trial stimulation
12 Hz, 0.5 s
Run onset
120 cm
n.s.
First-lick time (s)
Ctrl. Stim.
n.s.
First-lick distance (cm)
Ctrl. Stim.
n.s.
Reward rate (%)
Ctrl. Stim.

**Fig. S4 Speed controls and temporal specificity of LC stimulation effects, related to Fig. 1**

(**A**) Speed-matching procedure for early- versus late-first-lick trials. Left, mean running speed profiles before matching; middle, speed profiles after matching (early, median = 49.22 cm/s, IQR = [43.80, 52.08] cm/s; late, median = 47.33 cm/s, IQR = [43.20, 51.39] cm/s; p = 0.49; Wilcoxon signed-rank test). Right, comparison of run-onset acceleration after matching (early, median = 58.55 $cm/s^2$, IQR = [47.03, 63.33] $cm/s^2$; late, median = 57.38 $cm/s^2$, IQR = [43.37, 63.23] $cm/s^2$; p = 0.68; Wilcoxon signed-rank test). (**B**) Relationship between LC RON firing rate and mean running speed. Left, speed profiles for low- versus high-speed trial groups. Right, corresponding RON firing rate profiles aligned to run onset (run-onset FR: low speed, median = 2.97 Hz, IQR = [2.38, 3.59] Hz; high speed, median = 3.21 Hz, IQR = [2.82, 3.69] Hz; p = 0.17; 8 animals, 40 recordings; Wilcoxon signed-rank test). (**C**) First-lick distance for control versus phasic LC stimulation trials (control, median = 150.00 cm, IQR = [137.98, 162.31] cm; stimulation, median = 157.72 cm, IQR = [147.89, 162.86] cm; p = 2.97e-03; 12 animals, 54 recordings; Wilcoxon signed-rank test). (**D**) Phasic LC activation at reward. Left, schematic of reward-zone stimulation. Right, number of licks after reward delivery (control, median = 14.58, IQR = [8.05, 16.83]; stimulation, median = 14.29, IQR = [8.73, 17.51]; p = 0.43; 4 animals, 10 recordings; Wilcoxon signed-rank test). (**E**) Phasic LC activation delivered at mid-trial (first-lick time: control, median = 2.89 s, IQR = [2.75, 2.97] s; stimulation, median = 2.85 s, IQR = [2.65, 3.35] s; p = 0.70; first-lick distance: control, median = 162.00 cm, IQR = [152.10, 165.40] cm; stimulation, median = 164.70 cm, IQR = [154.90, 169.90] cm; p = 0.41; proportion of trials rewarded: control, median = 1.00, IQR = [1.00, 1.00]; stimulation, median = 0.97, IQR = [0.90, 1.00]; p = 0.06; 4 animals, 11 recordings; all Wilcoxon signed-rank tests).

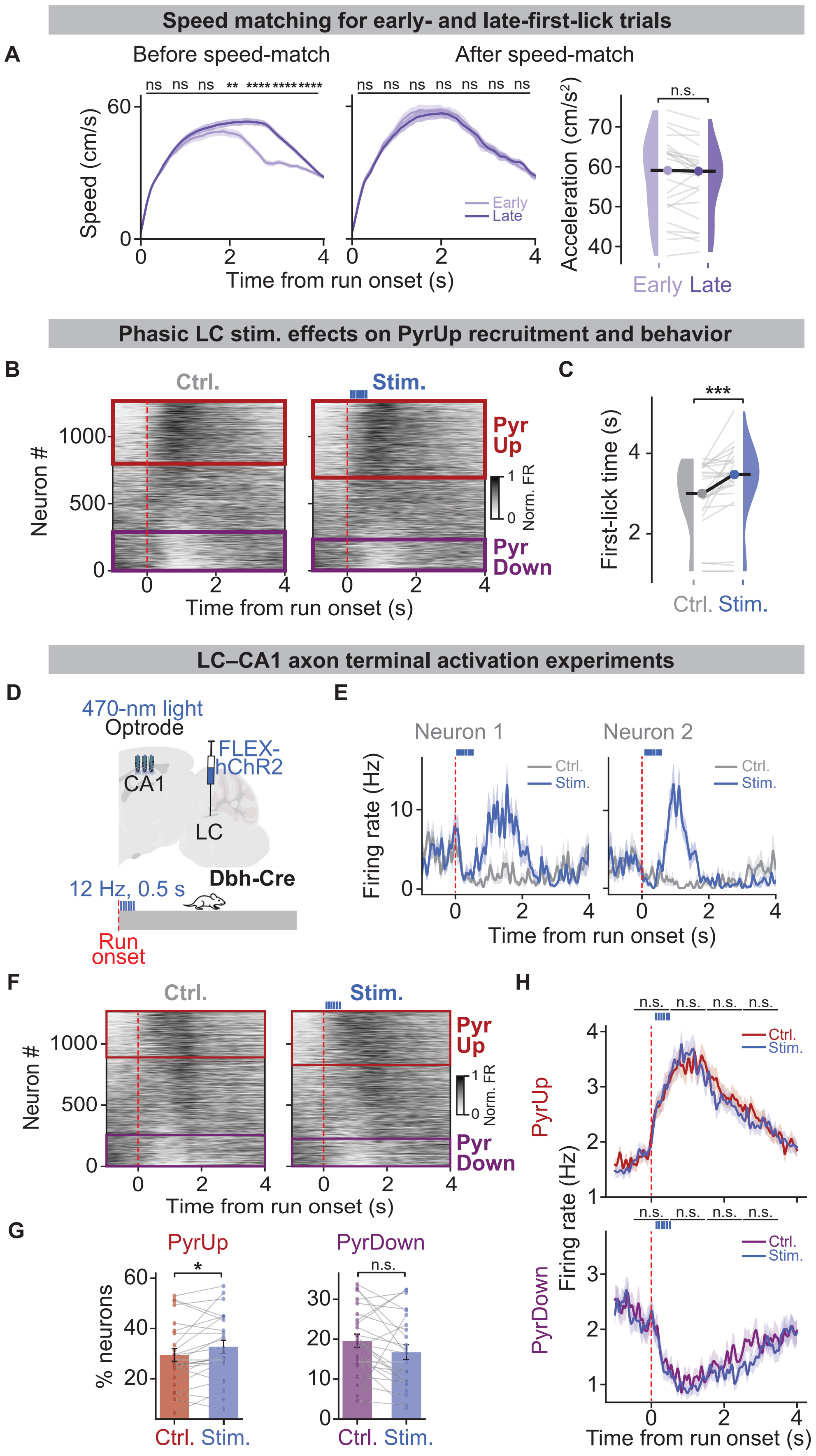

Speed matching for early- and late-first-lick trials
A
Before speed-match
After speed-match
ns ns ns ** ************
ns ns ns ns ns ns ns
Speed (cm/s)
Time from run onset (s)
Early
Late
n.s.
Acceleration (cm/s²)
Early Late
Phasic LC stim. effects on PyrUp recruitment and behavior
B
Ctrl.
Stim.
Neuron #
Pyr Up
Pyr Down
Norm. FR
Time from run onset (s)
C
***
First-lick time (s)
Ctrl. Stim.
LC–CA1 axon terminal activation experiments
D
470-nm light
Optrode
FLEX-hChR2
CA1
LC
Dbh-Cre
12 Hz, 0.5 s
Run onset
E
Neuron 1
Neuron 2
Ctrl.
Stim.
Firing rate (Hz)
Time from run onset (s)
F
Ctrl.
Stim.
Neuron #
Pyr Up
Pyr Down
Norm. FR
Time from run onset (s)
G
PyrUp
PyrDown
*
n.s.
% neurons
Ctrl. Stim.
Ctrl. Stim.
H
n.s. n.s. n.s. n.s.
PyrUp
PyrDown
Ctrl.
Stim.
Firing rate (Hz)
Time from run onset (s)

**Fig. S5 Speed-matching controls and comparison of LC somatic versus axon terminal stimulation effects on CA1 dynamics, related to Fig. 2**

(**A**) Running speed profiles for early- and late-first-lick trials before (left) and after (middle) speed-matching for the CA1 recordings shown in **Fig. 2C** (early, median = 43.85 cm/s, IQR = [41.78, 47.26] cm/s; late, median = 45.11 cm/s, IQR = [42.12, 46.92] cm/s; p = 0.15; Wilcoxon signed-rank test). Right, comparing run-onset acceleration after matching (early, median = 59.10 cm/s$^2$, IQR = [50.40, 63.87] cm/s$^2$; late, median = 58.88 cm/s$^2$, IQR = [51.08, 61.02] cm/s$^2$; p = 0.11; 17 animals, 25 recordings; Wilcoxon signed-rank test). (**B**) CA1 population heatmaps of trial-averaged firing rates aligned to run onset for control (left) and phasic LC stimulation (right) trials (statistics in **Fig. 2F**). Red square: PyrUp; purple square: PyrDown. Neurons are ordered by run-onset response index R within each condition. (**C**) First-lick timing for control (left) and phasic LC stimulation (right) trials (control, median = 2.94 s, IQR = [2.49, 3.61] s; stimulation, median = 3.42 s, IQR = [3.04, 3.75] s; p = 3.01e-04; 4 animals, 27 recordings; Wilcoxon signed-rank test). (**D-H**) CA1 silicon-probe recordings combined with optogenetic stimulation of LC–CA1 axon terminals. (**D**) Schematic of experiment. (**E**) Two example CA1 pyramidal neurons, showing trial-averaged firing-rate traces aligned to run onset for interleaved stimulation (blue) and control (gray) trials. Red dashed line indicates run onset. (**F**) CA1 population heatmaps, same layout as (**B**). (**G**) Across recordings, the percentage of PyrUp (left) and PyrDown neurons (right) in control and stimulation trials. Each dot represents one recording (control vs stimulation: PyrUp, 29.49 ± 2.54% vs 32.74 ± 2.58%; p = 0.021; PyrDown, 19.61 ± 1.64% vs 16.74 ± 1.83%; p = 0.076; 5 animals, 25 recordings; both paired *t*-tests). (**H**) Mean firing rate profiles for PyrUp (top) and PyrDown (bottom) neurons that maintained their classification in both control and stimulation trials (independent *t*-test for each time bin).

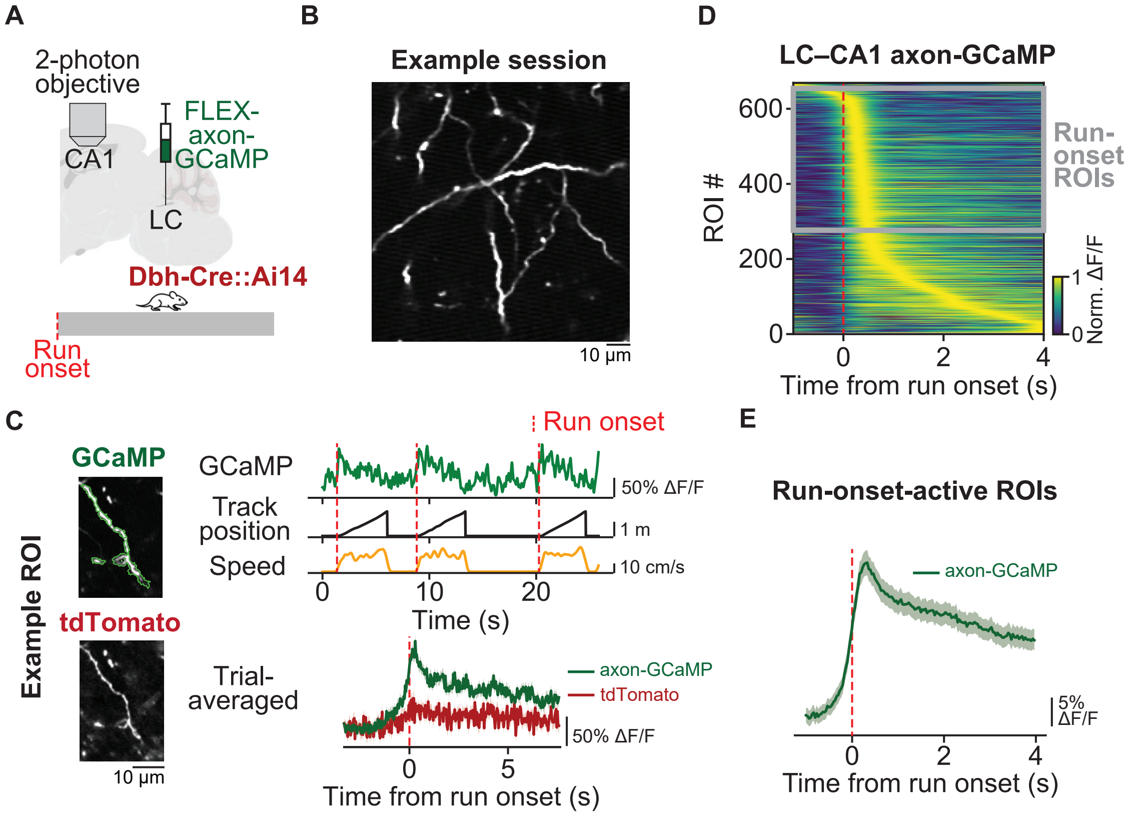
A
2-photon objective
CA1
FLEX-axon-GCaMP
LC
Dbh-Cre::Ai14
Run onset
B
Example session
10 μm
D
LC–CA1 axon-GCaMP
ROI #
600
400
200
0
Run-onset ROIs
Norm. ΔF/F
1
0
0
2
4
Time from run onset (s)
C
Example ROI
GCaMP
tdTomato
10 μm
Run onset
GCaMP
50% ΔF/F
Track position
1 m
Speed
10 cm/s
0
10
20
Time (s)
Trial-averaged
axon-GCaMP
tdTomato
50% ΔF/F
0
5
Time from run onset (s)
E
Run-onset-active ROIs
axon-GCaMP
5% ΔF/F
0
2
4
Time from run onset (s)

**Fig. S6 LC–CA1 Dbh$^+$ axons show run-onset calcium transients, related to Fig. 2**

(**A**) Experimental schematic of two-photon imaging of LC–CA1 axonal GCaMP activity. (**B**) Example field showing axon-GCaMP-labeled LC axons in CA1 from one recording. (**C**) Example axonal ROI. Left, axon-GCaMP channel (top) and the corresponding tdTomato reference channel (bottom). Right top, axon-GCaMP activity during consecutive trials (green) with corresponding track position (black) and running speed (orange). Right bottom, trial-averaged axon-GCaMP (green) and tdTomato (red) signals aligned to run onset. Red dashed lines indicate run onset. (**D**) Population heatmap of axon-GCaMP activity aligned to run onset (red dashed line) across LC–CA1 axonal ROIs. ROIs are ordered by activity peak time. The gray square highlights run-onset ROIs. (**E**) Mean signal traces of run-onset active ROIs.

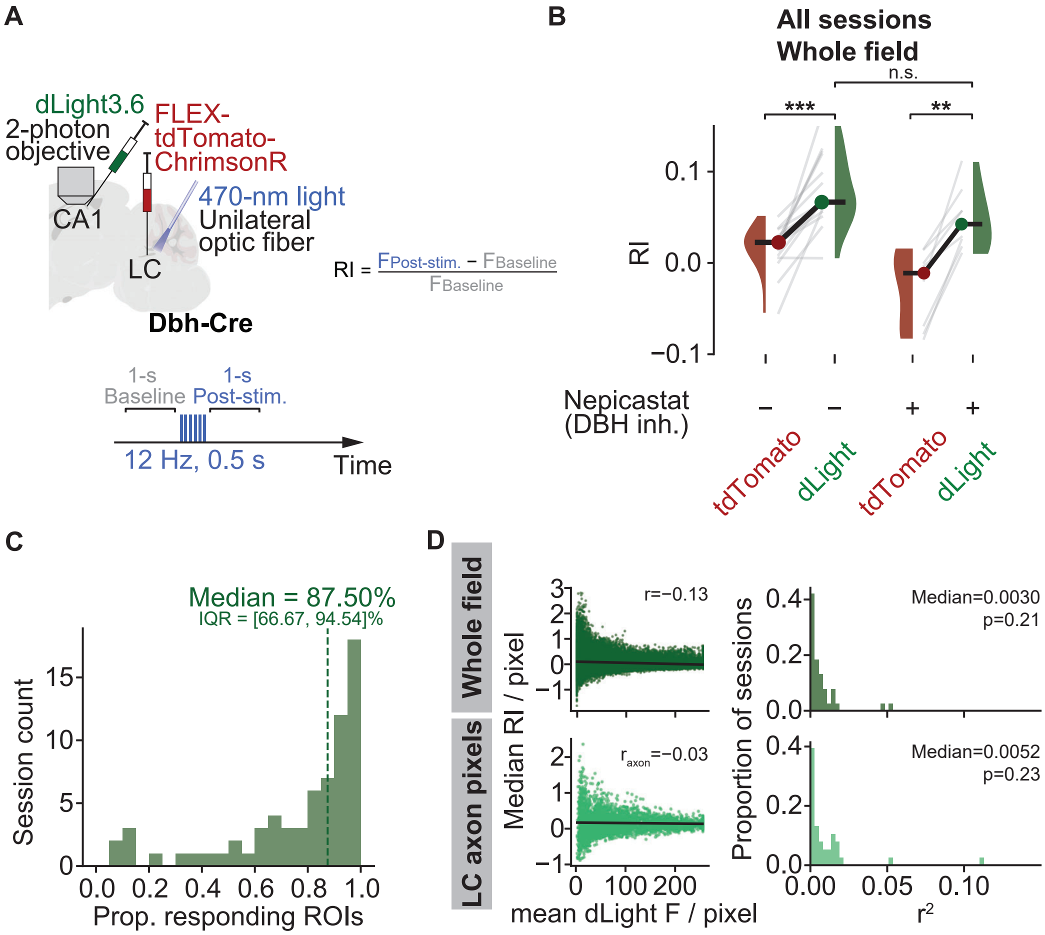
A
dLight3.6
2-photon objective
FLEX-tdTomato-ChrimsonR
CA1
470-nm light
Unilateral optic fiber
LC
Dbh-Cre
RI = FPost-stim. − FBaseline / FBaseline
1-s Baseline
1-s Post-stim.
12 Hz, 0.5 s
Time
B
All sessions Whole field
n.s.
***
**
RI
Nepicastat (DBH inh.)
tdTomato
dLight
C
Median = 87.50%
IQR = [66.67, 94.54]%
Session count
Prop. responding ROIs
D
Whole field
LC axon pixels
r=−0.13
raxon=−0.03
Median RI / pixel
mean dLight F / pixel
Median=0.0030
p=0.21
Median=0.0052
p=0.23
Proportion of sessions
r2

**Fig. S7 Validation of LC-evoked dLight signals: dopamine specificity and expression-level controls, related to Fig. 3**

(**A**) Schematic for CA1 dLight recordings with LC stimulation. Inset shows the same calculation of response index as in **Fig. 3C**. (**B**) RI measured in the tdTomato reference channel (red) and dLight channel (green) during recording sessions without (−) or with (+) DBH inhibitor nepicastat injection. Gray lines connect paired values within a recording (control: tdTomato RI, median = 0.02, IQR = [0.01, 0.03]; dLight RI, median = 0.07, IQR = [0.06, 0.11]; p = 1.22e−04. nepicastat: tdTomato RI, median = −0.01, IQR = [−0.07, 0.00]; dLight RI, median = 0.04, IQR = [0.02, 0.06]; p = 7.81e−03; both Wilcoxon signed-rank tests. dLight RI: control vs nepicastat, p = 0.071; 3 animals, 9 recordings; Wilcoxon rank-sum test). (**C**) Histogram across recordings of the proportion of LC-axon ROIs with significant dLight responses after stimulation. Unlike the pixel-wise analysis in **Fig. 3**, this analysis was performed at the level of individual LC-axon ROIs (see **Materials and methods**). (**D**) Relationship between pixelwise mean dLight fluorescence and pixelwise RI. Left, example recording showing pixelwise scatter and linear fit for the full field of view (top) and for LC-axon pixels only (bottom); r denotes Pearson's correlation coefficient. Right, histogram of the coefficient of determination ($r^2$) across recordings for the full field of view (top) and for LC-axon pixels only (bottom) (both one-sample *t*-tests).

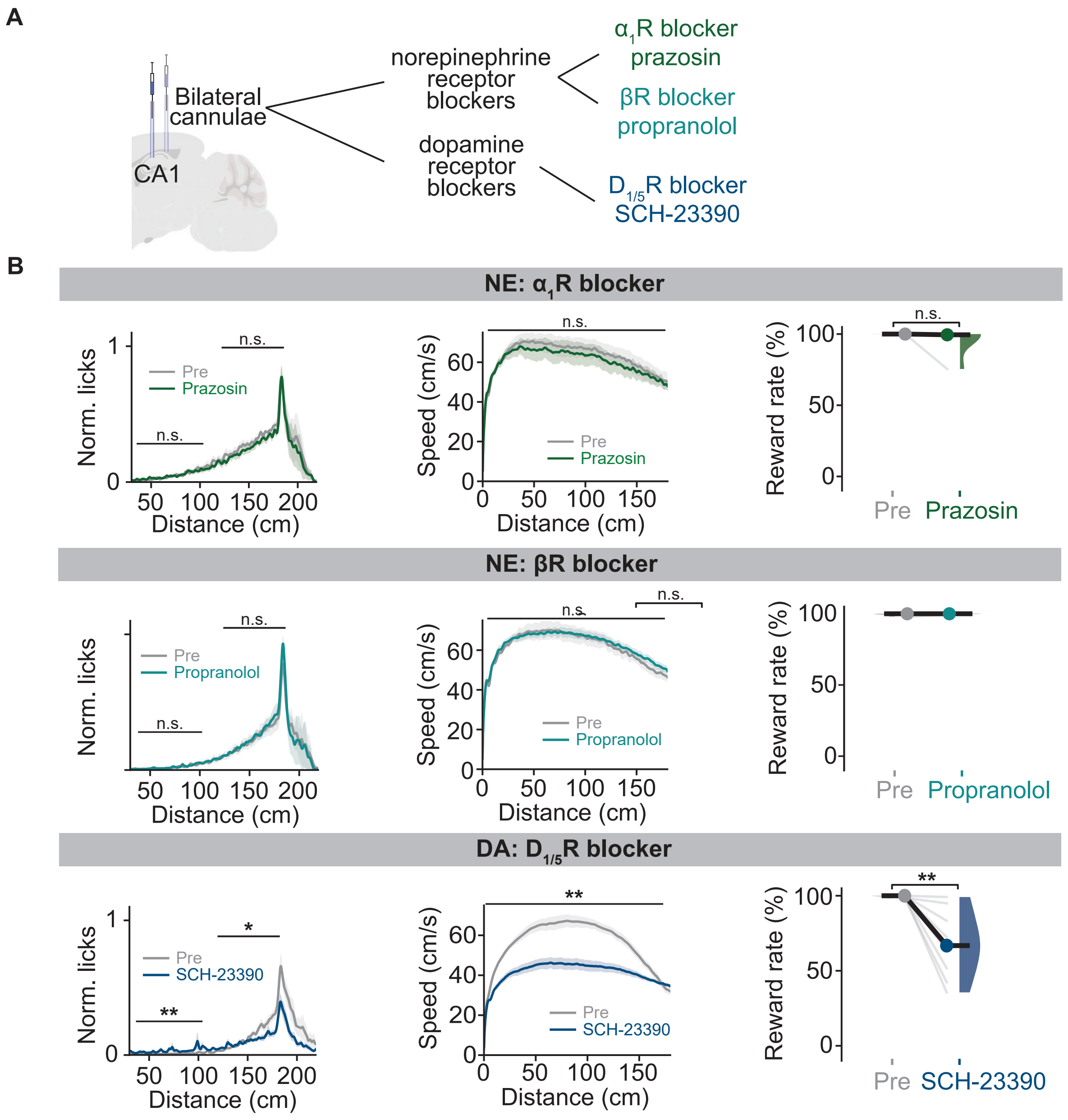

A
Bilateral cannulae
CA1
norepinephrine receptor blockers
dopamine receptor blockers
α1R blocker prazosin
βR blocker propranolol
D1/5R blocker SCH-23390
B
NE: α1R blocker
NE: βR blocker
DA: D1/5R blocker
Norm. licks
Speed (cm/s)
Reward rate (%)
Distance (cm)
Pre
Prazosin
Propranolol
SCH-23390
n.s.
**
*

**Fig. S8  Local pharmacological blockade of NE and DA receptors in CA1, related to Fig. 3**

(**A**) Schematic of bilateral receptor antagonist infusion into CA1. (**B**) Licking profiles (left), speed as a function of distance (middle) and reward rate (right) for pre versus drug infusion sessions (prazosin, anticipatory licking [120, 180]: pre, median = 0.27, IQR = [0.24, 0.32]; drug, median = 0.26, IQR = [0.23, 0.30]; p = 0.44; licks [30, 100] cm: pre, median = 0.046, IQR = [0.021, 0.050]; drug, median = 0.047, IQR = [0.029, 0.060]; p = 1.00; speed: pre, median = 57.30 cm/s, IQR = [50.29, 59.78] cm/s; drug, median = 50.52 cm/s, IQR = [47.56, 54.31] cm/s; p = 0.44; reward rate: pre, median = 100%, IQR = [100, 100]%; drug, median = 99%, IQR = [98, 100]%; p = 0.11; 3 animals, 6 recordings; propranolol, anticipatory licking [120, 180]: pre, median = 0.23, IQR = [0.22, 0.24]; drug, median = 0.21, IQR = [0.20, 0.22]; p = 0.81; licks [30, 100] cm: pre, median = 0.016, IQR = [0.006, 0.030]; drug, median = 0.017, IQR = [0.011, 0.017]; p = 0.63; speed: pre, median = 48.52 cm/s, IQR = [48.17, 58.86] cm/s; drug, median = 51.49 cm/s, IQR = [50.48, 51.64] cm/s; p = 1.00; reward rate: pre, median = 100%, IQR = [99, 100]%; drug, median = 100%, IQR = [100, 100]%; p = 0.65; 3 animals, 5 recordings; SCH-23390, anticipatory licking [120, 180]: pre, median = 0.14, IQR = [0.11, 0.17]; drug, median = 0.099, IQR = [0.077, 0.11]; p = 2.73e-02; licks [30, 100] cm: pre, median = 0.008, IQR = [0.004, 0.012]; drug, median = 0.030, IQR = [0.026, 0.033]; p = 1.95e-03; speed: pre, median = 48.56 cm/s, IQR = [45.48, 50.01] cm/s; drug, median = 36.97 cm/s, IQR = [34.89, 38.90] cm/s; p = 1.95e-03; reward rate: pre, median = 100%, IQR = [100, 100]%; drug, median = 67%, IQR = [55, 81]%; p = 1.95e-03; 4 animals, 10 recordings; all Wilcoxon signed-rank tests).

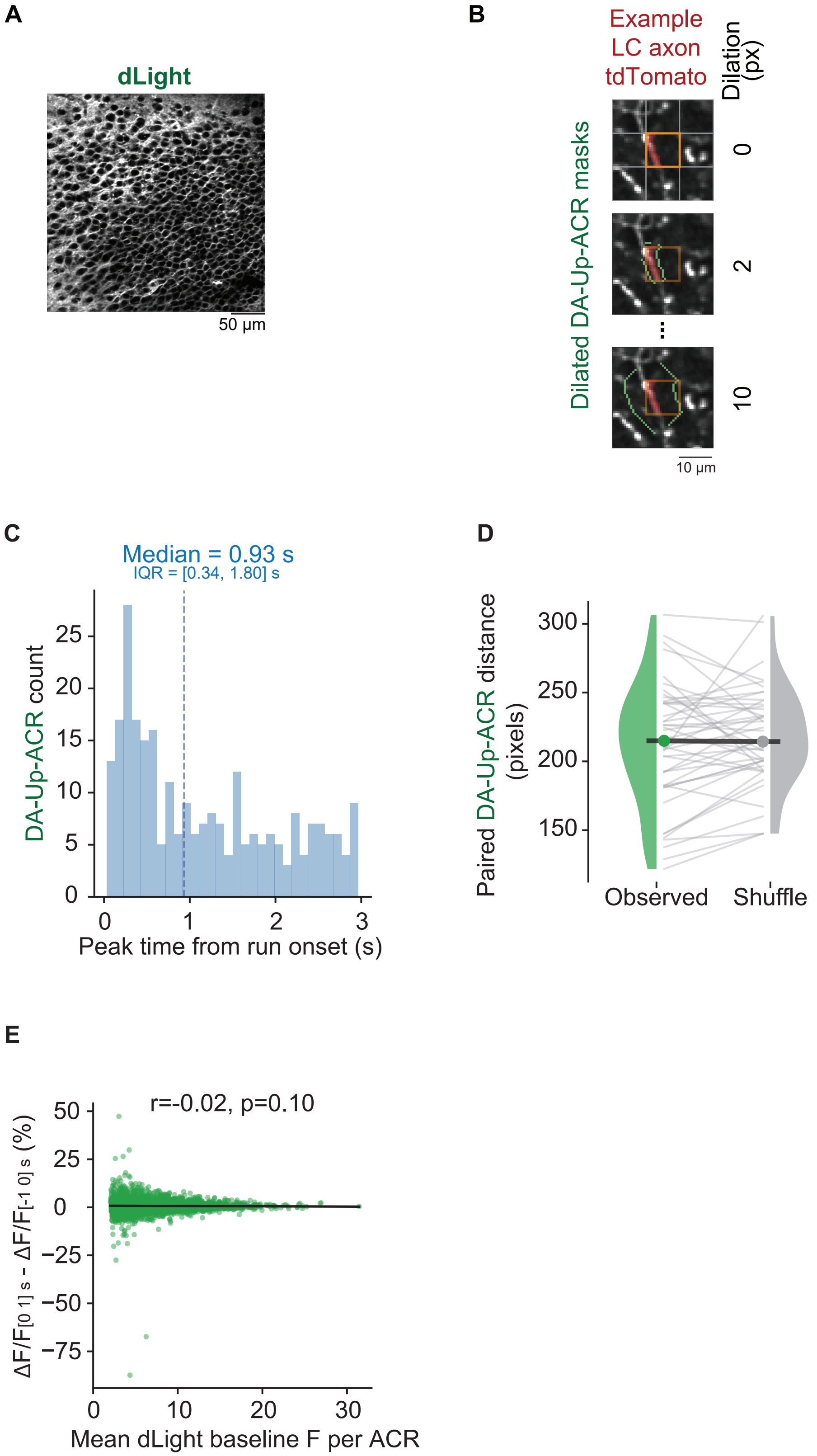
A
dLight
50 µm
B
Example
LC axon
tdTomato
Dilation
(px)
Dilated DA-Up-ACR masks
0
2
10
10 µm
C
Median = 0.93 s
IQR = [0.34, 1.80] s
DA-Up-ACR count
25
20
15
10
5
0
0
1
2
3
Peak time from run onset (s)
D
Paired DA-Up-ACR distance
(pixels)
300
250
200
150
Observed
Shuffle
E
r=-0.02, p=0.10
ΔF/F[0 1] s - ΔF/F[-1 0] s (%)
50
25
0
−25
−50
−75
0
10
20
30
Mean dLight baseline F per ACR

**Fig. S9  Timing and expression-level control of run-onset dLight transients near LC axons in CA1, related to Fig. 3**

(**A**) Example field of view expressing dLight3.6. (**B**) Example spatial dilation masks (green rings) of a DA-Up-ACR. (**C**) Histogram of peak times (relative to run onset) for DA-Up-ACRs. Peak time was measured from the trial-averaged trace for each ACR. Dashed line marks the median (median = 0.93 s, IQR = [0.34, 1.80] s). (**D**) Pairwise distances between DA-Up-ACRs in each recording, compared with pairwise distances obtained after randomly shuffling DA-Up labels among all ACRs (paired *t*-test for median distance of each session, p = 0.25; 4 animals, 46 recordings). **(E)** Relationship between the change in dLight signal for each ACR, computed as ΔF/F in the [0, 1] s window minus ΔF/F in the [−1, 0] s window around run onset, and the mean dLight baseline F. r denotes Pearson's correlation coefficient.

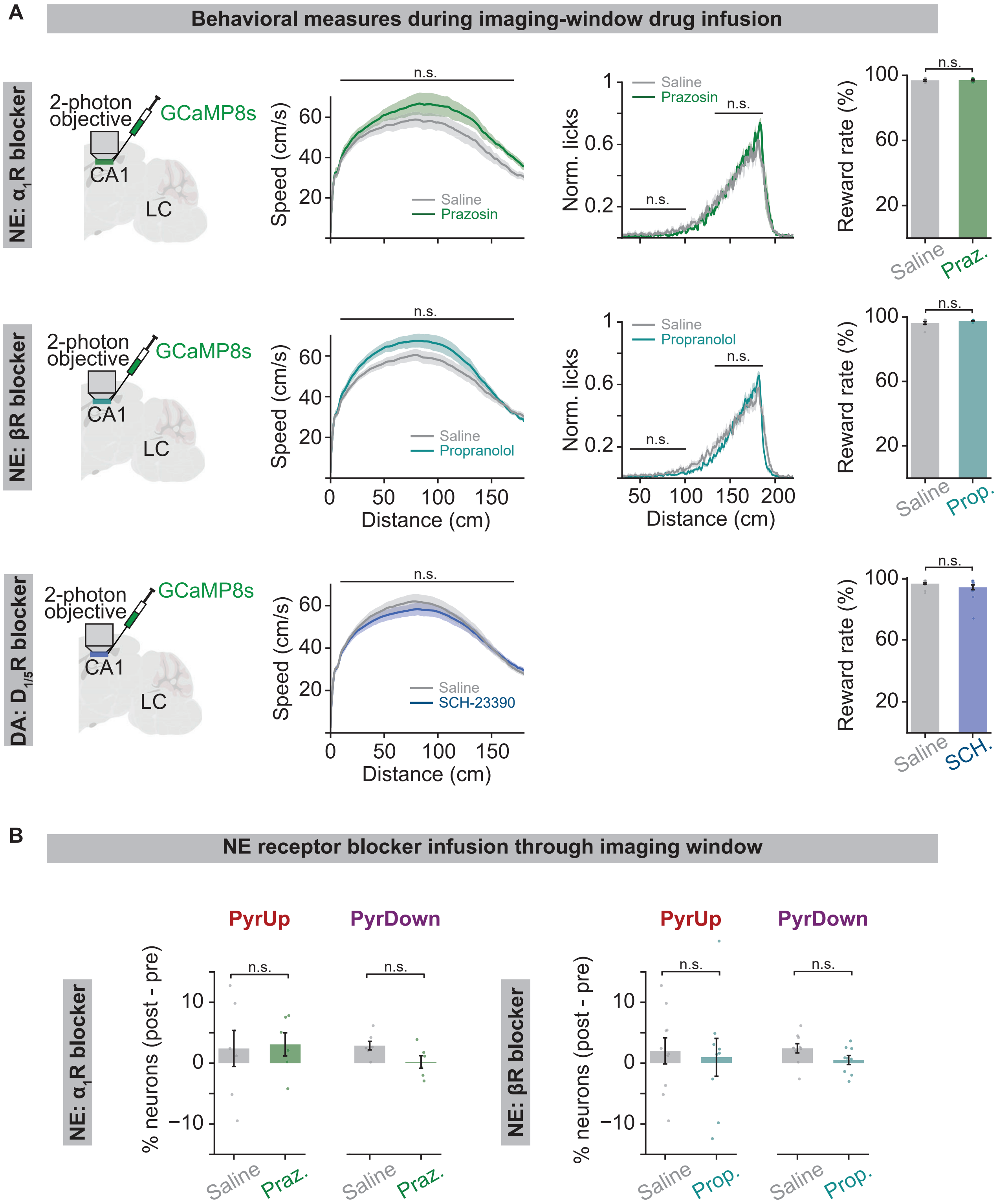
A
Behavioral measures during imaging-window drug infusion
NE: α1R blocker
2-photon objective
GCaMP8s
CA1
LC
Speed (cm/s)
n.s.
Saline
Prazosin
Norm. licks
Reward rate (%)
Saline
Praz.
NE: βR blocker
Propranolol
Prop.
Distance (cm)
DA: D1/5R blocker
SCH-23390
SCH.
B
NE receptor blocker infusion through imaging window
PyrUp
PyrDown
% neurons (post - pre)
NE: α1R blocker
NE: βR blocker

**Fig. S10 Local pharmacological blockade of NE and DA receptors in CA1 during two-photon imaging, related to Fig. 4**

(**A**) Behavioral measures during two-photon imaging after local infusion of receptor antagonists: $\alpha_1$-adrenergic antagonist prazosin (top row), β-adrenergic antagonist propranolol (middle row), and $D_{1/5}R$ antagonist SCH-23390 (bottom row). Left column, experimental schematics. Second column: mean running speed. Third column: licking profiles, normalized to the maximum lick rate across saline and drug conditions. Right column: reward rate. Saline: gray. Drug: colored traces as indicated in each panel (prazosin, speed: saline, median = 47.31 cm/s, IQR = [44.62, 50.84] cm/s; drug, median = 53.54 cm/s, IQR = [49.50, 57.54] cm/s; p = 0.09; licks [30, 100] cm: saline, median = 0.017, IQR = [0.011, 0.020]; drug, median = 0.018, IQR = [0.010, 0.024]; p = 0.78; anticipatory licking [120, 180] cm: saline, median = 0.334, IQR = [0.270, 0.406]; drug, median = 0.354, IQR = [0.331, 0.370]; p = 0.48; reward rate: saline, median = 97.12%, IQR = [95.92, 98.10]%; drug, median = 97.63%, IQR = [95.97, 98.12]%; p = 0.62; 3 animals, 7 recordings for saline, 6 recordings for prazosin; propranolol, speed: saline, median = 48.36 cm/s, IQR = [46.24, 51.12] cm/s; drug, median = 54.78 cm/s, IQR = [50.27, 60.38] cm/s; p = 0.14; licks [30, 100] cm: saline, median = 0.016, IQR = [0.009, 0.020]; drug, median = 0.011, IQR = [0.007, 0.024]; p = 0.62; anticipatory licking [120, 180] cm: saline, median = 0.300, IQR = [0.256, 0.339]; drug, median = 0.280, IQR = [0.262, 0.352]; p = 0.87; reward rate: saline, median = 96.82%, IQR = [95.44, 98.11]%; drug, median = 98.06%, IQR = [97.22, 98.11]%; p = 0.25; 4 animals, 10 recordings for saline, 9 recordings for propranolol; SCH-23390, speed: saline, median = 49.42 cm/s, IQR = [44.02, 55.36] cm/s; drug, median = 47.38 cm/s, IQR = [41.21, 53.67] cm/s; p = 0.60; reward rate: saline, median = 97.06%, IQR = [95.66, 97.67]%; drug, median = 96.64%, IQR = [91.80, 97.41]%; p = 0.44; 8 animals, 19 recordings for saline, 16 recordings for SCH-23390; all Wilcoxon rank-sum tests). (**B**) Across recordings, change in percentage of PyrUp and PyrDown neurons compared to pre-infusion sessions after saline (gray) or drug (left, prazosin (praz.); right, propranolol (prop.)) infusion experiments (left, percentage of PyrUp change: saline, 2.41 ± 2.97%; prazosin, 3.09 ± 1.90%; p = 0.85; percentage of PyrDown change: saline, 2.88 ± 0.71%; prazosin, 0.19 ± 1.04%; p = 0.06; 3 animals, 7 recordings for saline, 6 recordings for prazosin; right, percentage of PyrUp change: saline, 2.02 ± 2.15%; propranolol, 0.96 ± 3.11%; p = 0.78; percentage of PyrDown change: saline, 2.44 ± 0.76%; propranolol, 0.51 ± 0.75%; p = 0.09; 4 animals, 10 recordings for saline, 9 recordings for propranolol; all independent *t*-tests).

A

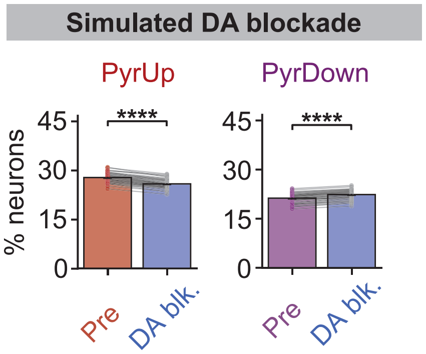
Simulated DA blockade
PyrUp
PyrDown
% neurons
45
30
15
0
****
Pre
DA blk.

**Fig. S11  Simulated DA blockade in the LC–dopamine–CA1 model, related to Fig. 5**

(**A**) Percentage of PyrUp (left) and PyrDown (right) neurons in the model before and after simulated DA blockade. Gray lines connect paired results in individual simulation runs (PyrUp, pre, median = 28.30%, IQR = [27.20, 29.32]%; blockade, median = 26.20%, IQR = [25.35, 27.30]%; p = 7.49e-10; PyrDown, pre, median = 21.80%, IQR = [20.92, 22.60]%; blockade, median = 23.05%, IQR = [21.90, 23.80]%; p = 7.31e-10; both Wilcoxon signed-rank tests).
